# Reorganization of the Flagellum Scaffolding Induces a Sperm Standstill During Fertilization

**DOI:** 10.1101/2023.06.22.546073

**Authors:** Martina Jabloñski, Guillermina M. Luque, Matías D. Gómez-Elías, Claudia Sanchez-Cardenas, Xinran Xu, Jose Luis de la Vega-Beltran, Gabriel Corkidi, Alejandro Linares, Victor X. Abonza Amaro, Aquetzalli Arenas-Hernandez, María Del Pilar Ramos-Godinez, Alejandro López-Saavedra, Dario Krapf, Diego Krapf, Alberto Darszon, Adan Guerrero, Mariano G. Buffone

## Abstract

Mammalian sperm delve into the female reproductive tract to fertilize the female gamete. The available information about how sperm regulate their motility during the final journey to the fertilization site is extremely limited. In this work, we investigated the structural and functional changes in the sperm flagellum after AE and during the interaction with the eggs. The evidence demonstrates that the double helix actin network surrounding the mitochondrial sheath of the midpiece undergoes structural changes prior to the motility cessation. This structural modification is accompanied by a decrease in diameter of the midpiece and is driven by intracellular calcium changes that occur concomitant with a reorganization of the actin helicoidal cortex. Midpiece contraction occurs in a subset of cells that undergo AE, live-cell imaging during in vitro fertilization showed that the midpiece contraction is required for motility cessation after fusion is initiated. These findings provide the first evidence of the F-actin network’s role in regulating sperm motility, adapting its function to meet specific cellular requirements during fertilization, and highlighting the broader significance of understanding sperm motility.

**Significant statement:** In this work, we demonstrate that the helical structure of polymerized actin in the flagellum undergoes a rearrangement at the time of sperm-egg fusion. This process is driven by intracellular calcium and promotes a decrease in the sperm midpiece diameter as well as the arrest in motility, which is observed after the fusion process is initiated.

## Introduction

Sperm motility is required for arrival at the site of fertilization and to penetrate the different layers surrounding the egg. The temporal regulation of sperm motility involves the concerted action of multiple cell structures to enable fertilization. The initial motility of ejaculated sperm is characterized by a linear progressive movement as the cells traverse a long distance within the female reproductive tract. However, at one point, before reaching the oocyte, this initial progressive motion must be changed to a vigorous non-progressive mode of motility called hyperactivation (*1*). During that migration, most of mouse sperm undergo acrosomal exocytosis (AE), which takes place in the upper segments of the oviduct, before the sperm directly interact with the egg or its surrounding layers (*2–4*). This exocytic event is critical because proteins involved in this process are rearranged in preparation for fusion(*5*, *6*). A second dramatic change in motility is subsequently required for efficient sperm-egg fusion. During this event, which has not been studied in detail so far, sperm completely stop moving (*7–9*). The cease in sperm motility is considered as an indicative marker of an effective fusion between gametes, given that it is necessary to complete the attachment and fusion between sperm and eggs. Fusion itself is a complex event mediated by several proteins identified using loss of function strategies (*10*, *11*). Nevertheless, the available information about how sperm regulate their motility during the final journey to the fertilization site and during the interaction with the female gamete is extremely limited. In this final journey, most sperm cells migrate to the ovulated eggs after AE, and very little is known about the motility state of those sperm and about the molecular mechanism in charge of the motility arrest.

The regulation of sperm motility is achieved by balancing force transduction and the mechanical properties of the flagellum. Both effects are governed by the flagellum cytoskeleton, which consists of two major components: a microtubule-based axoneme, located at the flagellum axis, and actin filaments around it (*12*). The midpiece and the principal piece also contain the outer dense fibers and the fibrous sheath that have traditionally been referred to as part of the sperm cytoskeleton. Recently it was found that the three-dimensional organization of polymerized actin in the flagellum midpiece of murine sperm forms a unique double helix arrangement accompanying mitochondria (*13*). This spatial distribution does not extend into the principal piece, where actin is uniformly distributed between the axoneme and the plasma membrane. Across all investigated cells, this double helix consisted of exactly 87 gyres with a pitch of 244±1 nm, yielding a total midpiece length in mice of 21.2±0.3 μm (*13*). Such accurate control of sizes is not typical in live cells given that reactions are prone to stochastic effects and entropy leads to large cell-to-cell fluctuations. Thus, the precise control of the actin helix comes at a substantial regulatory cost for sperm cells. Nevertheless, the function of this specialized structure of actin filaments is largely unknown.

In this work, we find definite evidence for the role of the helical actin structure in the final stage of motility regulation, when sperm dramatically shift from being hyperactivated to a practically immotile stage. We have investigated the structural changes in the actin cytoskeleton of the sperm flagellum after AE and during the interaction with the eggs. By using a combination of single-cell imaging and super-resolution microscopy methods, our results demonstrate that the helical structure of polymerized actin undergoes a radical structural change at the time of sperm-egg fusion. Further, we uncover that this process is triggered by a substantial calcium ([Ca^2+^]_i_) influx. This actin-dependent signaling pathway promotes a decrease of the sperm midpiece diameter and motility arrest, both of which are found to be required to complete the fusion process during fertilization.

## Results

### AE promotes a cessation of motility in a subset of sperm cells

After AE, sperm need to accomplish migration to the ampulla, reach unfertilized eggs, penetrate the cumulus matrix and the zona pellucida and finally, undergo fusion with the oocyte (*14*). All these processes share the need to modulate sperm motility. Little is known about the regulation of motility after the occurrence of AE or prior to the fusion with the oocyte.

To simultaneously monitor the acrosomal status and sperm motility in live cells, transgenic mice whose sperm express enhanced green fluorescent protein (EGFP) in the acrosome and red fluorescent protein (DsRed2) in the mitochondria were used (Figure 1A, Supplementary movie S1). EGFP-DsRed2 sperm were immobilized at the head on laminin-coated coverslips, while still allowing free flagellar movement (Figure 1B, Supplementary movie S2). While the presence of the acrosome is monitored using the EGFP fluorescence signal, motility was assessed through the movement of the midpiece (DsRed2 fluorescence), which was beating in and out of the imaging plane (Figure 1B).

**Figure 1.**
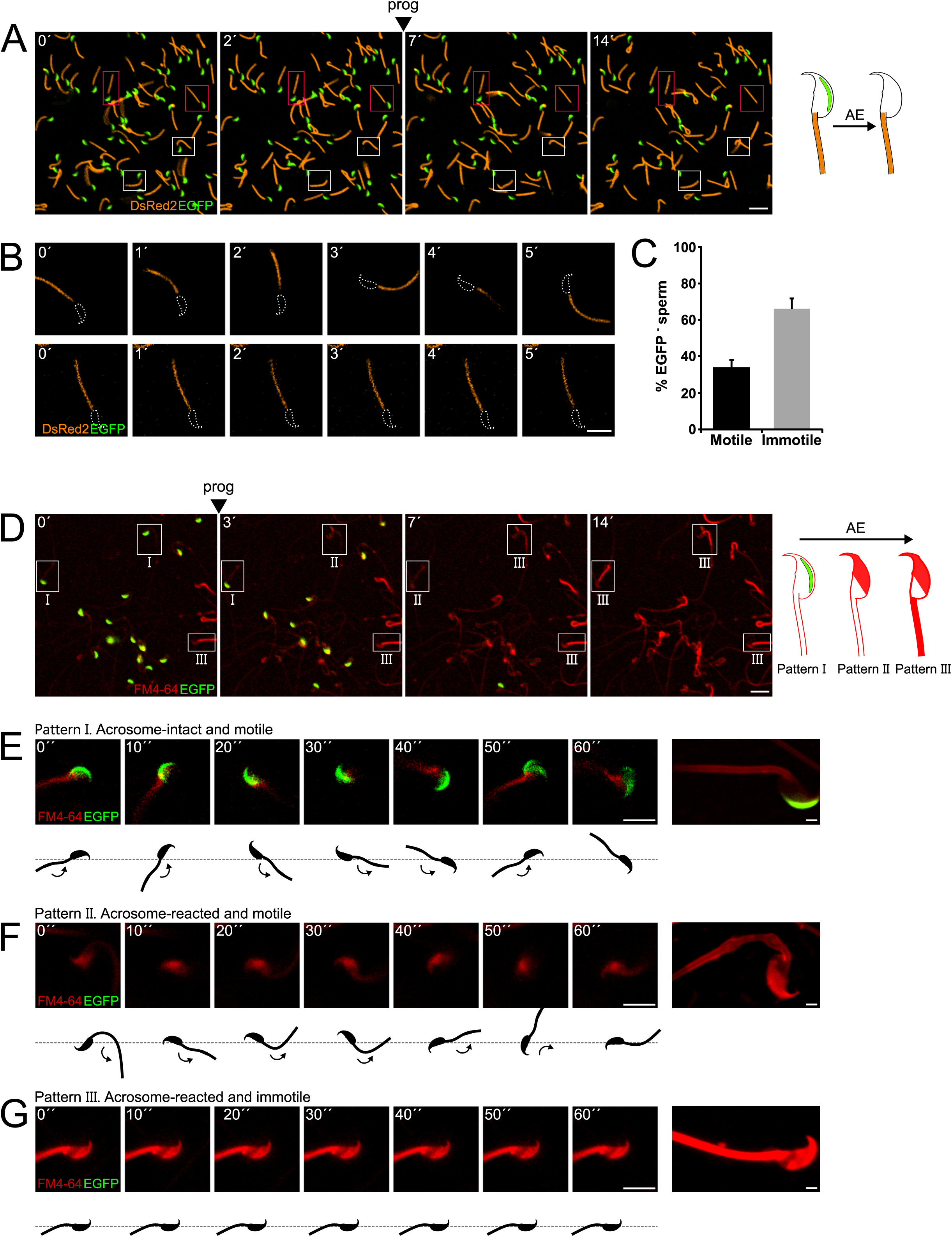
Sperm motility loss and FM4-64 fluorescence dynamics in acrosome-reacted transgenic EGFP-DsRed2 sperm. A) Representative time series of transgenic EGFP-DsRed2 sperm attached to concanavalin A-coated coverslips, with AE induced by 100 μM progesterone. White squares indicate cells with spontaneous AE (prior to induction), while pink squares highlight cells with progesterone-induced AE. A schematic representation of AE in this transgenic model is shown on the right side of the panel. Scale bar = 20 μm. B) Representative time series of transgenic EGFP-DsRed2 sperm that have already experienced AE, attached to laminin-coated coverslips. The upper panel displays a cell with motility after AE, and the lower panel shows an immotile cell. DsRed2 is presented in orange, and EGFP in green. Scale bar = 10 μm. C) Quantification of motile and immotile acrosome-reacted sperm (EGFP-). A total of 235 cells were counted across at least three independent experiments. D) Representative time series of transgenic EGFP-DsRed2 sperm stained with 10 μM FM4-64 and attached to concanavalin A-coated coverslips, with AE induced by 100 μM progesterone. White squares indicate cells exhibiting patterns I, II, or III after progesterone induction. Scale bar = 20 μm. A schematic representation of AE in this transgenic model stained with FM4-64 is shown on the right side of the panel. E-G) Representative images of capacitated transgenic EGFP-DsRed2 sperm stained with 10 μM FM4-64. Panel E) displays an acrosome-intact, motile sperm (Pattern I), F) shows an acrosome-reacted sperm with motility and low FM4-64 midpiece fluorescence (Pattern II), and G) presents an acrosome-reacted sperm with no motility and high FM4-64 midpiece fluorescence (Pattern III). In all three cases, cells were induced with 100 μM progesterone. Scale bar = 10 μm. Enlarged images of each pattern are shown on the right panel. Scale bar = 2 μm. Representative images from at least five independent experiments are displayed.

The majority of acrosome-intact sperm were able to move. Interestingly, Figures 1B and 1C show the coexistence of two populations of sperm that underwent AE (cells lacking EGFP signal) induced upon addition of progesterone, a physiological trigger of AE. Some of the acrosome reacted sperm moved normally (34.1±3.7 %, n = 235), whereas the majority of them remained immotile (65.9 ± 6.2 %, n = 2350) (Figures 1B and 1C, upper and lower panel, respectively, Supplementary movie S2).

The flagellar beat cycle encompasses a self-regulatory mechanism that receives feedback from molecular and mechanical signals (*15*). The complete abortion of flagellar beat cycle observed in Figures 1B and 1C might be indicative of a stimulus provided by or occurring concomitant with AE. To comprehend the connection between AE and motility, we hypothesized the existence of a mechanical change in the flagella as a result of the AE. For this reason, AE was studied in the presence of FM4-64 fluorescent dye (*16*) to visualize structural changes at the plasma membrane, which can occur at macroscopic scales observed as an alteration in the shape of the flagellum, or at mesoscopic scales, monitored by local changes in FM4-64 fluorescence occurring at the vicinity or within the plasma membrane. EGFP-DsRed2 sperm were stimulated with progesterone and their ‘motility’ behavior was recorded for 5 min in the presence of FM4-64. The loss of EGFP fluorescence in the acrosome of transgenic mice correlated with a noticeable increase of FM4-64 fluorescence in the head, as shown in Figure 1D (see panels 0’ and 3’, Supplementary movie S3). In addition, we noticed that sperm that underwent AE, later showed an increase in the FM4-64 fluorescence intensity in the flagellum as well (see panel 14’of Figure 1D, Supplementary movie S3).

A more detailed analysis of individual cells demonstrated changes in FM4-64 fluorescence according to their motility (Supplementary movie S4). Figure 1E shows a motile acrosome-intact sperm, (a control case), where the fluorescence levels of EGFP and FM4-64 remained constant during the entire experiment (Pattern I). Interestingly, sperm that lost their acrosome (no EGFP and high FM4-64 fluorescence in the sperm head) and continued moving, presented low levels of FM4-64 fluorescence in the midpiece (Figure 1F, Pattern II). On the other hand, acrosome-reacted sperm that remained immotile showed a significant rise in the FM4-64 midpiece fluorescence (Figure 1G, Pattern III). Noteworthy, both subsets of cells (with and without motility) displayed the expected increase in FM4-64 fluorescence in the sperm head.

To deepen our understanding on how sperm lose motility during AE, we designed a sperm tracking system that can also monitor changes in fluorescence in moving cells (Figure S1B and S1C). Three parameters were then assessed in real time, namely, the beat frequency of the flagellum (by tracking it on bright field images, or the DsRed2 channel, Figure S1A), the status of the acrosome (EGFP signal), and changes occurring at or within the membrane in the midpiece (FM4-64 signal). Figure S1B shows that sperm that did not undergo AE remained motile with a stable beat frequency and low FM4-64 fluorescence over the recording time. In stark contrast, after AE, cells gradually diminished the beating frequency until a complete arrest is observed, while a gradual increase in FM4-64 fluorescence in the midpiece is observed (Figure S1C).

To evaluate the relationship between between FM4-64 and viability, a vital dye Sytox Blue was used in imaging flow cytometry experiments. Fluorescence intensities in non-capacitated sperm stimulated with ionomycin (Figure S1D) were determined and two populations with Sytox Blue signals were clearly distinguished (Sytox+ and Sytox-), enabling the discernment between live and dead sperm. Interestingly, the upper right panels (Sytox Blue+ / FM4-64 high) consistently show a positive correlation between FM4-64 and Sytox Blue. Nonetheless, the lower panels (Sytox Blue-) show no correlation with FM4-64 fluorescence, indicating that this population can exhibit either low or high FM4-64 fluorescence. Single-cell examples are shown, where the four categories are represented: dead sperm with low FM4-64 fluorescence (Sytox Blue + / FM4-64 low), dead sperm with high FM4-64 fluorescence (Sytox Blue + / FM4-64 high), live sperm with low FM4-64 fluorescence (Sytox Blue - / FM4-64 low), and live sperm with high FM4-64 fluorescence (Sytox Blue - / FM4-64 high). Therefore, while the FM4-64 signal alone is not a definitive marker for either AE or cell death, it is crucial to use additional viability assessments, such as Sytox Blue, to accurately differentiate between live and dead sperm in studies of acrosome exocytosis and sperm motility. Cell viability was always considered, as any imaged sperm was chosen based on motility, indicated by a beating flagellum. The determination of whether selected sperm die during or after AE remains to be elucidated. The results presented in Figure 1 and Supplementary Figure S1 show examples of motile sperm that experience an increase in FM4-64 fluorescence.

Altogether, these experiments demonstrate that AE promotes a cease of motility in a subset of sperm cells, which coincides with an increase in FM4-64 fluorescence in the midpiece.

### AE is followed by a decrease in the midpiece diameter

Live-cell super-resolution microscopy was used to understand the structural changes occurring in the midpiece. Figure 2A shows a wide-field image of a sperm midpiece stained with FM4-64 (left) and analyzed using super-resolution radial fluctuations (SRRF, right) (*17*). AE dynamics was monitored by FM4-64 fluorescence in the head (Figure 2B, upper right insets). A decrease in the midpiece diameter over time was observed following AE either being spontaneous, induced by a physiological agonist (progesterone), or induced by a non-physiological surge of Ca^2+^ (ionomycin) (Figures 2B and 2C and Supplementary movie S5). In the negative control (no AE), the diameter remained unchanged. This phenomenon was also observed using mean shift super-resolution (MSSR) (*18*). To further support this observation, other plasma membrane probes of different molecular structure, such as Memglow 700 (Figure S2A), Bodipy-GM1 (Figure S2B), and FM1-43 (Figure S2C), were used. In all three cases, a decrease in the midpiece diameter was observed after AE.

**Figure 2.**
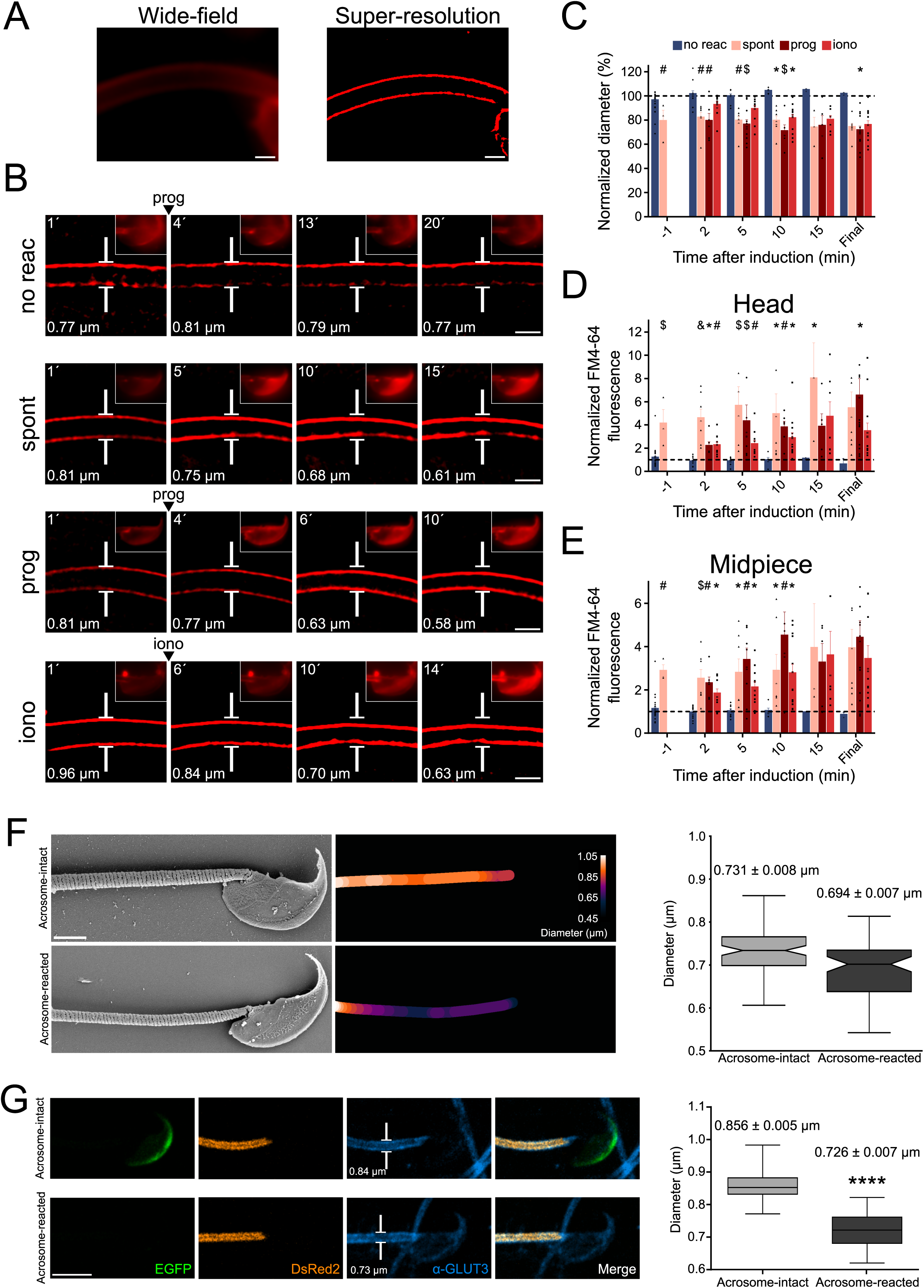
Midpiece contraction coincides with the onset of AE. A) Left panel displays a wide-field fluorescence image of capacitated CD1 sperm membrane stained with 0.5 μM FM4-64, while the right panel shows its super-resolution SRRF reconstruction. Scale bar = 1 μm. B) Representative time series of sperm midpiece with no AE (no reac), spontaneous exocytosis (spont), progesterone (prog, 100 μM) and ionomycin-induced (iono, 10 μM) exocytosis, respectively. Following acquisition, images were analyzed using SRRF. Insets in the sperm head show wide-field images of AE. The midpiece diameter value is displayed in the bottom left corner for each time point. Scale bar = 1 μm. C) Quantification of midpiece diameter changes for each experimental group across time. Data are presented as a percentage of the initial diameter value before induction for each cell. D-E) Quantification of FM4-64 fluorescence in the sperm head and midpiece, respectively, for each experimental group across time. Data are presented as times of increases compared to initial fluorescence before AE induction. *p<0.05; #p<0.01; $p<0.001 and &p<0.0001 compared to the non-reacted group. A nonparametric Kruskal-Wallis test was performed in combination with Dunn’s multiple comparisons test. Representative images of at least 5 independent experiments are shown, with 36 cells analyzed. F) Comparison of the midpiece architecture in acrosome-intact (AI, upper panel) and acrosome-reacted (AR, lower panel) sperm using scanning electron microscopy (SEM). Representatives images are shown, middle panels show quantification of these images whereas the left panel shows the quantification of all replicates. Data is presented as mean ± SEM, Kruskal-Wallis test was employed, p = 0.013, (AI n=85, AR n=72). Scale bar = 2 μm. G) Capacitated transgenic EGFP-DsRed2 sperm were induced by 100 µM progesterone. Cells were fixed and immunostained against -GLUT3 in order to see the plasma membrane in the midpiece. Representative images of at least 2 independent experiments are shown. Left panel shows quantification of midpiece diameter in acrosome-intact and acrosome-reacted EGFP-DsRed2 sperm. Data is presented as mean ± sem. A nonparametric Mann Whitney test was performed, ****p<0.001 (AI n=84, AR n=47). Scale bar = 5 μm.

The FM4-64 fluorescence intensity in the head and in the midpiece (Figures 2D and 2E) were also assessed. As expected, the increase in FM4-64 fluorescence in the head occurs after AE. Furthermore, in agreement to what was observed in Figure 1, fluorescence intensity in the midpiece also increased over time in sperm that underwent AE (Figures 2D and 2E).

To confirm these results, two alternative methods to visualize the change in sperm midpiece diameter were used. In neither of them, a membrane dye was used. First, indirect immunofluorescence to detect a membrane protein (GLUT3) was performed. As shown in Figure 2G, a decrease in the midpiece diameter was also observed. Second, scanning electron microscopy (SEM) was used to evaluate the midpiece in acrosome-intact or reacted sperm (Figure 2F). The overall diameter of the midpiece in acrosome-intact sperm was larger than in acrosome-reacted sperm, with measurements of 0.731 ± 0.008 μm and 0.694 ± 0.007 μm, respectively . Overall, we have confirmed using three different approaches that AE is followed by a midpiece diameter.

### The contraction of the midpiece initiates in the proximal part of the flagellum

To investigate whether the contraction of the midpiece is triggered at a random location or in a particular region of the flagellum, kymographs were used. A kymograph allows visualization of dynamical aspects of a given phenomenon in a single figure, where the temporal dimension is expressed as an axis of the image through a spatial-temporal transformation of the dataset. Two types of kymographs were used (Figure 3A). First, super-resolution microscopy kymographs were built to monitor dynamic changes in the diameter of the flagellum along the midpiece. Second, to investigate the number of contraction sites, fluorescence kymographs were made from diffraction-limited images to observe changes in FM4-64 fluorescence over time and across the entire midpiece. Interestingly, Figure S2D shows a significant negative correlation between the midpiece diameter and the FM4-64 fluorescence, unveiling a tool to obtain an approximate value of the midpiece diameter without the need of super-resolution microscopy.

**Figure 3.**
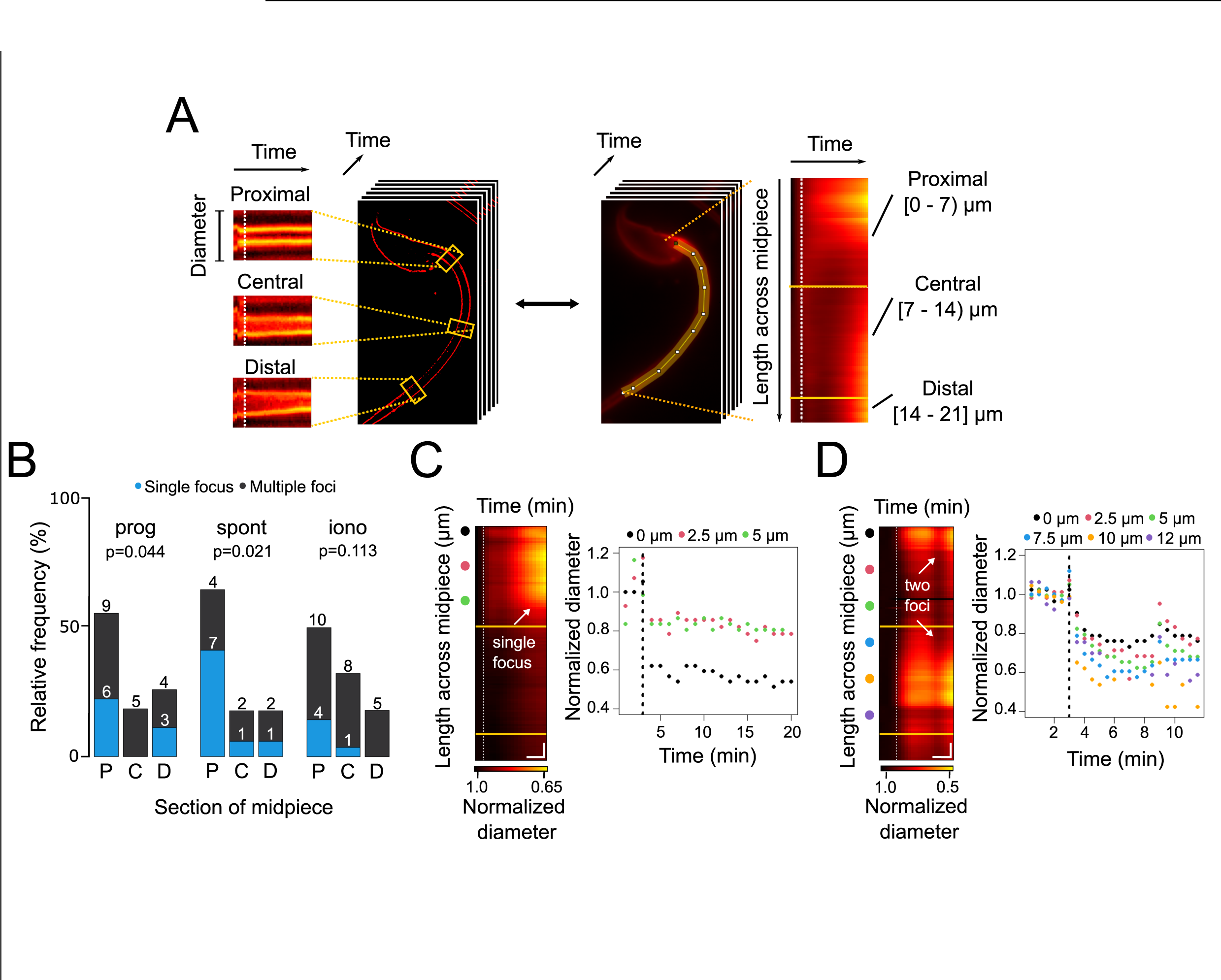
Contraction initiation preferentially occurs near the head-midpiece junction. A) Schematic diagram illustrating the generation of super-resolution kymographs from SRRF-processed images. Crosslines are drawn every 2.5 μm through the sperm midpiece, and the Image-J Kymograph builder plug-in is used to create kymographs. The x-axis represents time, and the y-axis shows diameter changes. For wide-field images, a line along the midpiece is drawn to create fluorescence kymographs, with the y-axis representing midpiece length. Three sections of the midpiece are defined: proximal [0-7 μm], central [7-14 μm], and distal [14-21 μm]. B) Relative frequency graph displaying the distribution of the initiation sites for midpiece contractions in sperm with no AE (no reac), spontaneous exocytosis (spont), progesterone-induced (prog, 100 μM) exocytosis, and ionomycin-induced (iono, 10 μM) exocytosis, respectively. The x-axis indicates the midpiece section where the contraction begins: proximal (P), central (C), or distal (D). A chi^2^ test was performed using the R language environment. C-D) Representative contraction kymographs and diameter measurements for progesterone-induced (100 μM) AE with one or two contraction initiation sites, respectively. In contraction kymographs, yellow lines demarcate midpiece sections, and colored spots indicate where super-resolution kymographs were created. Both kymograph and diameter measurement graphs display a dotted vertical line marking the induction point. For C, horizontal scale bar = 5 min and vertical scale bar = 1 μm and for D, horizontal scale bar = 3 min and vertical scale bar = 1 μm. Data from at least 5 independent experiments are shown, with 36 cells analyzed.

Three experimental subsets that underwent AE were assessed: spontaneous, induced by progesterone, and induced by ionomycin. To scrutinize randomness of focus-driven contraction, the midpiece was segmented in three regions (Figure 3A): proximal (near the neck), central, and distal (near the annulus). In these groups, it was then evaluated whether cells presented one or more foci of contraction.

Figure 3B shows a cumulative study, which indicates that contraction is preferentially initiated at the proximal part of the midpiece, regardless of being progesterone-induced or spontaneous AE. When ionomycin was used as an agonist of AE, the contraction began randomly in any segment of the midpiece. Figures 3C and 3D show two examples of flagellar dynamics for progesterone-induced sperm, which either presented one or two foci of contraction. In both cases, the diameter decreased along the whole midpiece, seen as a transition from dark to bright red colors in Figures 3C and 3D. Noteworthy, focal points of contraction were consistently observed: i.e., a case with a single focus (Figure 3C), or a case with two foci of contraction (Figure 3D). Figures S2E and S2F show ionomycin-induced sperm that present one or more foci. It is observed in the normalized diameter kymograph that the flagellum presents a single focus between the measurements of 7.5 µm and 10 µm (Figure S2E, left panel). This is reflected in the right panel in the light blue and orange dots, which are the actual diameter measurements that decreased first and most (Figure S2E). A representative behavior within this experimental group is represented in Figure S2F, where the entire midpiece contracted (all its sections simultaneously). This is observed in the color scale of the normalized diameter kymograph and coincides with the real measurements of the diameter in the right panel, which decreased their value. Lastly, Figures S2G and S2H show examples of spontaneous AE with one or two foci of contraction, respectively. In particular, in Figure S2H, a first focus of contraction between 0 and 5 µm that coincided with the diameter measurements of the right panel that decreased their value first and to a greater extent (black, pink and green dots). Although the entire midpiece contracted, a second focus of contraction appeared around the measurements of 10 to 15 µm and coincidentally, these are the points with the strongest decrease in the diameter (orange, purple and red points).

### The reduction of the midpiece diameter occurs concomitantly with a [Ca^2+^]_i_ increase in the flagellum

It is widely accepted that an increase in [Ca^2+^]_i_ precedes AE (*16*, *19*). It has been shown that AE initiation by progesterone occurs after an increase in [Ca^2+^]_i_ in the sperm head that is later propagated towards the midpiece (*19*). To comprehend the connection between AE and the concomitant contraction of the midpiece, we hypothesized the existence of a signal, i.e., Ca^2+^, which propagates from the head to the flagellum and modulate the architecture of the sperm flagellum.

To investigate if the increase in the concentration of [Ca^2+^]_i_ in the midpiece is correlated with the midpiece contraction, sperm were incubated with FM4-64 and Fluo4, a Ca^2+^ fluorescent sensor (Supplementary movie S6). Figure 4A shows a live-cell imaging experiment of sperm cells that did not undergo AE (control case), nor a significant rise in [Ca^2+^]_i_. Noteworthy, upon induction with progesterone (Figure 4C, Supplementary movie S6), there was an increase in the midpiece [Ca^2+^]_i_ followed by an increase in FM4-64 fluorescence, which is indicative of a contraction of the flagellum (see Figure S2D). These single cell experiments illustrate a reduction of the midpiece diameter that occurs concomitantly with an increase in [Ca^2+^]_i_ within the flagellum. A large population of sperm was analyzed using low magnification (10X). Figures 4B and 4D show a collection of single cell kymographs, where each row represents the fluorescence of Fluo4 (green) and FM4-64 (red) of the midpiece over time. Unstimulated cells are displayed on Figure 4B and sperm that were exposed to progesterone are shown in Figure 4D. Most of the cells display a transient [Ca^2+^]_i_ increase followed by an increase in FM4-64 fluorescence. This pattern was also observed in ionomycin-induced cells (Figures S3A and S3B and Supplementary movie S6) with the exception that, as expected, in this case the rise in [Ca^2+^]_I_ was faster and sustained. These findings indicate that an [Ca^2+^]_i_ transient increase, happening in the midpiece, precedes the contraction of the flagellum.

**Figure 4.**
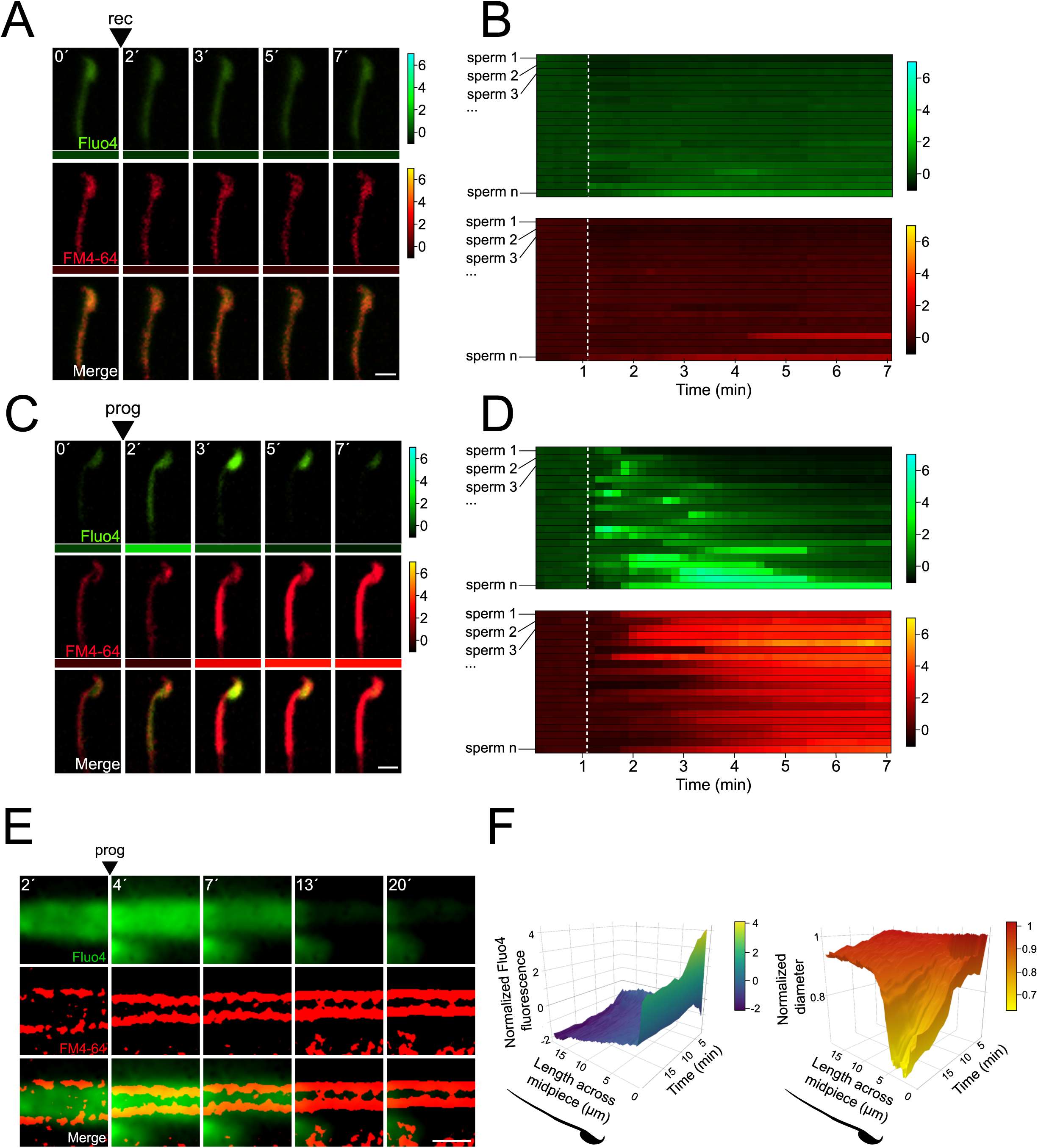
Midpiece contraction is driven by [Ca^2+^]_i_ changes. A) The representative time series demonstrates [Ca^2+^]_i_ and AE dynamics. Capacitated F1 sperm, loaded with Fluo4 AM, were immobilized on concanavalin A-coated coverslips and incubated in a recording medium (rec) containing 10 μM FM4-64. Rec was added as indicated by arrowheads. Scale bar = 10 μm. Beneath each frame in the Fluo4 (green) and FM4-64 (red) images, a color code displays the normalized intensity of the fluorescence signal (scale bar on the right of the panel). B) Kymograph-like analysis of the midpiece of 20 sperm following the addition of recording medium. Each row depicting the [Ca^2+^]_i_ (upper) and membrane (lower) dynamics of a single cell over time. A white dotted line indicates the moment of addition. The images presented are representative of at least five independent experiments. C) The representative time series demonstrates [Ca^2+^]_i_ and AE dynamics. Progesterone (prog, 100 μM) was added as indicated by arrowheads. Scale bar = 10 μm. Beneath each frame in the Fluo4 (green) and FM4-64 (red) images, a color code displays the normalized intensity of the fluorescence signal (scale bar on the right of the panel). D) Kymograph-like analysis of the midpiece of 20 sperm following the addition of prog. Each row depicting the [Ca^2+^]_i_ (upper) and membrane (lower) dynamics of a single cell over time. A white dotted line indicates the moment of addition. The images presented are representative of at least five independent experiments. Consistently, a [Ca^2+^]_i_ transient precedes contraction, which is proportional to the increase in FM4-64 fluorescence, as shown in Figure S2D. E) Representative time series of [Ca^2+^]_i_ and midpiece contraction dynamics. Capacitated CD1 sperm were loaded with Fluo4 AM, immobilized on concanavalin A-coated coverslips, and incubated in a recording medium containing 0.5 μM FM4-64. AE was induced with 100 μM progesterone (prog, arrowhead). Fluo4 images are widefield images, while FM4-64 images are SRRF-processed (super-resolution). Scale bar = 1 μm. F) 3D kymographs of [Ca^2+^]_i_ (left) and contraction (right) dynamics. Data are normalized to the mean of the frames before the induction of AE. Representative images from at least 5 independent experiments are shown, with 36 cells analyzed.

The relationship between the observed [Ca^2+^]_i_ rise and the flagellar contraction was then assessed using live-cell super-resolution imaging. Figure 4E shows there is a [Ca^2+^]_i_ increase in the midpiece that coincides in space and time with a contraction of the midpiece. To provide insight in the spatio-temporal relationship of the [Ca^2+^]_i_ rise and the midpiece contraction, these results were visualized using a 3D kymograph encompassing the dynamics happening along the entire midpiece (Figure 4F). Overall, a rise in [Ca^2+^]_i_ occurred along the whole midpiece, which coincided with a generalized contraction of the midpiece (Figure 4F). Remarkably, a transient focal increase of [Ca^2+^]_i_ was observed at the base of the flagellum, which correlates in space and time with a focal reduction of the diameter (Figure 4F). We then sought to identify a molecular/structural link between both processes.

### The distance between the actin cytoskeleton and the plasma membrane is decreased during the contraction of the midpiece

The midpiece is shaped by two major structural elements: the axoneme, which consists of tubulin and its accessory components, including the outer dense fibers (ODFs); and a recently described network of filamentous actin (F-actin) arranged in a helicoidal conformation along the midpiece (*13*). We investigated whether the actin network plays a role in regulating the midpiece contraction associated with AE. We considered three possible scenarios: (i) an unknown mechanical force, mediated by a rigid matrix such as F-actin, brings the plasma membrane closer to the center of the flagellum; (ii) a dynamic relationship exists between the plasma membrane and the actin cytoskeleton of the midpiece, facilitating contraction; and (iii) the midpiece contraction is driven by an uncharacterized signaling mechanism that acts independently of the actin cytoskeleton.

The role of the actin cytoskeleton in the midpiece contraction was investigated through live-cell super-resolution using SiR-actin, a fluorescent probe that binds to F-actin (*20–22*). We used FM4-64 to visualize the plasma membrane (Figure 5) and transgenic sperm expressing DsRed2 to observe the mitochondrial network (Figure S4). Two stimulated sperm, one that experienced AE (Figure 5B) and another that did not (Figure 5A) are shown. In both cases, a network of F-actin was observed beneath the plasma membrane of the midpiece (cortical actin), enveloping the mitochondrial network (Figure S4A). As expected, in the absence of AE, the midpiece diameter (observed through FM4-64) remained unchanged (see also Figure 2, and Figure S4). In this case, both the diameter of the F-actin network and its proximity to either the plasma membrane or the mitochondrial network remained static (Figure 5A), suggesting a structural role for the F-actin network within the midpiece (scenario - (i)), such as supporting the organization of the mitochondrial network (Figure S4). Remarkably, acrosome-reacted sperm (Figure 5B) experienced an abrupt structural reorganization of the midpiece, characterized by a decrease in the distance between the plasma membrane and both the F-actin and mitochondrial networks (Figures S4 and S5). This observation suggests that the remodeling of the midpiece structure is driven by a mechanical (or molecular) interaction occurring at the boundaries of the plasma membrane (supporting scenario - (ii)).

**Figure 5.**
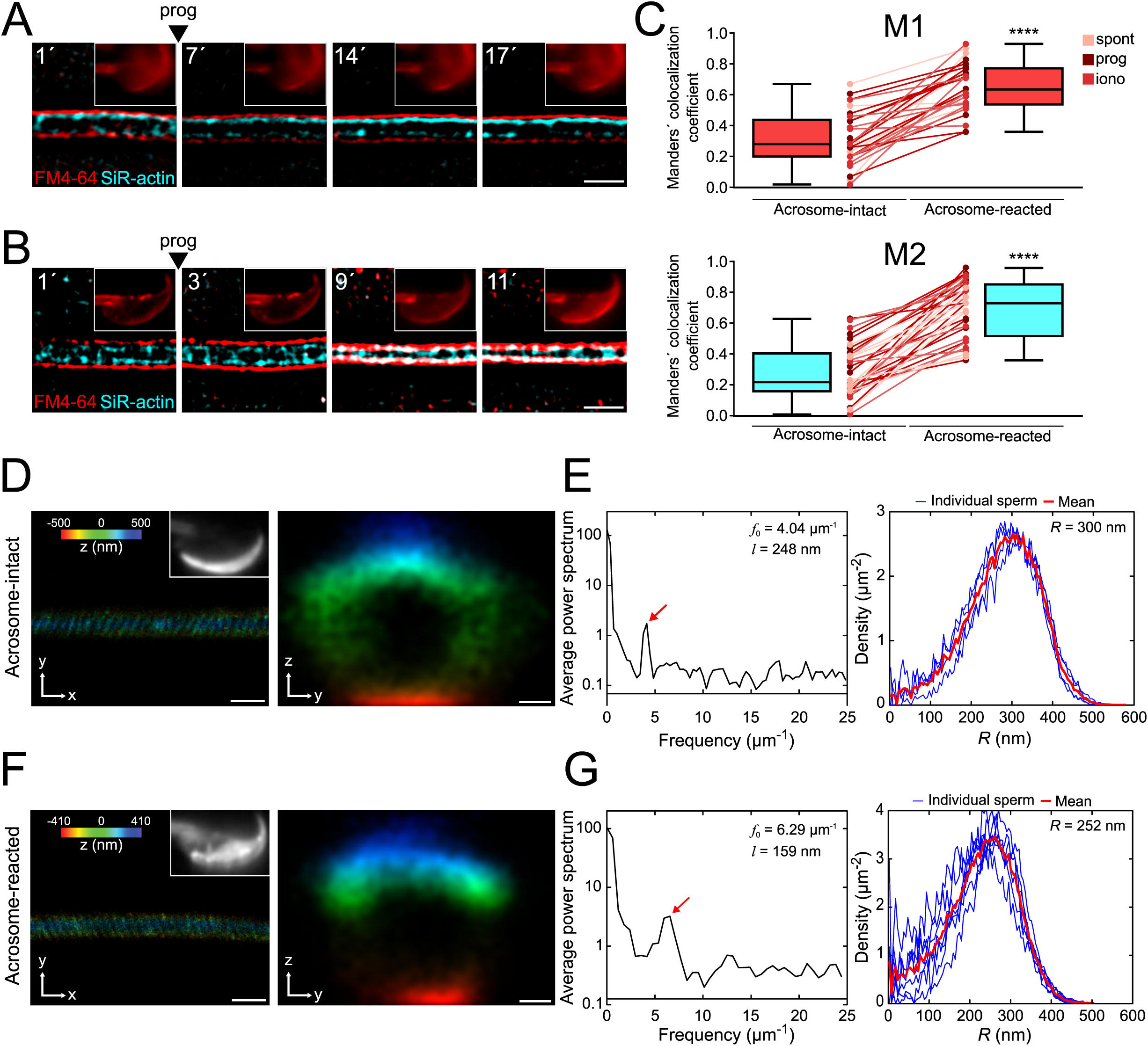
The flagellar membrane approaches the actin cytoskeleton in the midpiece of the sperm flagellum during midpiece contraction and AE. A-B) Representative time series of plasma membrane and actin cytoskeleton colocalization in the midpiece in the absence of AE (A) and during progesterone-induced AE (B, prog, 100 μM; FM4-64 shown in red, SiR-actin shown in cyan). Capacitated CD1 sperm were loaded with 100 nM SiR-actin, immobilized on concanavalin A-coated coverslips, and incubated in a recording medium containing 0.5 μM FM4-64. Scale bar = 1 μm. Representative images from at least 5 independent experiments are shown, with 36 cells analyzed. C) Manders’ colocalization coefficients for acrosome-intact and acrosome-reacted cells in the midpiece. M1 was assigned to FM4-64, and M2 to SiR-actin. Data are presented as mean ± SEM. ****p<0.0001 represents statistical significance. Paired t test was performed. D-G) Representative images of sperm midpiece stained with the acrosome marker PNA (left panel, upper right insets, epifluorescence) and phalloidin (actin filaments, STORM) for acrosome-intact (D-E) and acrosome-reacted (F-G) cells. The left panel displays a longitudinal section of the midpiece (Scale bar = 10 μm), while the right panel illustrates the radial distribution (Scale bar = 0.2 μm). E and G) Schematics of the analyzed actin double helix parameters in the midpiece: helical pitch (*l*, distance between turns of the helix, left panel), helical pitch frequency (*f*_0_, number of turns the helix makes per 1 μm), and radial distribution (R, radius of the double helix, right panel). Representative images from at least 3 independent experiments are shown. Four acrosome-intact cells and seven acrosome-reacted cells were analyzed.

To assess the potential interaction between the F-actin network and the plasma membrane (or its associated components), we examined their colocalization through live-cell super-resolution imaging. Figure 5C presents two Manders’ colocalization coefficients (*23*). M1 represents the proportion of F-actin (in pixels) that colocalizes with the plasma membrane, while M2 measures the proportion of plasma membrane pixels colocalizing with the F-actin network. A Manderś value of 1 indicates full colocalization, while a value of 0 implies no colocalization.

Figure 5C reveals that acrosome-intact sperm displayed low Manders’ coefficients (M1= 0.32 ± 0.03; M2= 0.28 ± 0.04). In contrast, sperm undergoing midpiece contraction exhibited a significant increase in colocalization between the F-actin network and the plasma membrane, as evidenced by the rise in M1 and M2 coefficients (M1= 0.65 ± 0.03; M2 = 0.69 ± 0.03). This increased colocalization occurred simultaneously with midpiece contraction.

Overall, these results confirm that (i) the contraction of the midpiece is linked to the remodeling of the plasma membrane and potentially the F-actin cytoskeleton, and that (ii) both structures interact with each other, directly or indirectly, at the nanoscales. We then seek to understand whether the driving force for the contraction of the midpiece emanates from the F-actin network. The following scenarios were envisaged: (a) the plasma membrane moves towards the actin cytoskeleton, (b) the actin cytoskeleton moves towards the plasma membrane, or (c) both structures come closer to the center of the flagellum. To investigate the occurrence of any of these scenarios, the positions of fluorescence peaks of FM4-64 and SiR-actin in super-resolution kymographs were tracked, as proxy of either membrane or actin cytoskeleton localization, respectively (Figures S5A and S5B). Figure S5C shows representative histograms of SiR-actin and FM4-64 over time. Both FM4-64 and SiR-actin fluorescence peaks came closer to each other. This effect can be also seen in Figure S5F, where acrosome-intact cells display a distance of 0.180 ± 0.007 μm between the plasma membrane and the actin cytoskeleton. In acrosome-reacted cells, the distance was as small as 0.074 ± 0.006 μm after midpiece contraction.

In Figure S5D, a section of the midpiece of an ionomycin-induced AE sperm is shown. In this case, the plasma membrane and the actin cytoskeleton approach move closer toward the center of the cell. (set to 0). The slope calculated from the linear fit denotes the velocity of this change. In this example, the plasma membrane is contracting at a rate of 14 μm/min, whereas the actin cytoskeleton is contracting at a rate of 3 μm/min (Figure S5E). In both cases, the signal presents a negative slope in sperm that underwent AE, consistent with a decrease in midpiece diameter. Since the results of the analysis of SiR-actin slopes were not conclusive, we studied the actin cytoskeleton structure in more detail.

### The actin cytoskeleton in the midpiece is remodeled during the AE

In the midpiece, polymerized actin forms a double helix that accompanies mitochondria (*13*). To investigate if the contraction of the midpiece is associated with structural modifications of the actin double-helix, 3D-STORM was used. Cells were exposed to progesterone, fixed, and co-stained with PNA and phalloidin to visualize the acrosomal status and the actin structure, respectively.

Figures 5D and 5F show representative images of both dyes for acrosome-intact and acrosome-reacted sperm (left panels). Different parameters were calculated: 1) Helical pitch (*l*), which is the distance between turns of the helix; 2) Frequency (*f*_0_ = 1/*l*), obtained from the Fourier transform of the image and represented the number of turns that the helix makes per unit length; and 3) Radial distribution (*R*), to infer the distance between the center of the midpiece and the maximum of fluorescence (*13*). The frequency increased substantially in the acrosome-reacted sperm compared to intact sperm (6.29 vs 4.04 μm^-1^, Figures 5E and 5G, red arrows in left panels). In addition, the helical pitch diminished its magnitude in acrosome-reacted sperm (159 nm vs 248 nm) (Figures 5E and 5G, left panels). The radial distribution of F-actin in cells that underwent AE indicated a smaller radius compared to acrosome-intact sperm (252 vs 300 nm, Figures 5E and 5G, right panel). Altogether, these results confirm that the actin double-helix undergoes structural changes during the contraction of the midpiece

To further investigate the structural rearrangements of the actin cytoskeleton during AE, a fluorescent molecule number and brightness analysis was performed (*24–26*). The number measure indicates the abundance or concentration of SiR-Actin molecules bound to F-actin fibers, while the brightness analysis helps reveal dynamic changes in molecular aggregation processes. This analysis is based on the idea that higher-order fluorescent complexes, which are mobile within the sample, will cause an increase in signal variability over time. An increase in brightness suggests the formation or movement of supramolecular structures, such as bundles of actin fibers bearing SiR-Actin. AE induction with progesterone and ionomycin resulted in a significant increase in actin number and brightness compared to the control (non-reacted) group (Figure S6A). Spontaneously reacted sperm also exhibited this behavior, albeit in a more moderate fashion (Figure S6A).

The observed increase in SiR-Actin number after AE induction suggests major actin monomer recruitment to the polymerizing fibers (Figure S6A and S6B), which produces an apparent increase in local fluorophore concentration (Figure S6B). This effect positively correlated with a similar increase in SiR-actin fluorescence (Figure S6A), which is known to indicate actin polymerization due to its high affinity for F-actin.

Actin cytoskeleton polymerization in the midpiece results in a signal variation of increased magnitude (an increase in SiR-Actin brightness) as it undergoes structural rearrangements (Figures S6A and S6B), which are compatible with other observations reported in this work. Both assembly and transport of actin complexes into higher oligomeric states (F-actin) across the cellular milieu led to an apparent increase in the registered SiR-Actin brightness (Figure S6B). Additionally, actin filament displacement along the mitochondrial sheath during AE provides a SiR-Actin brightness measure with high dispersion, indicating a redistribution of this cellular structure. These results suggest that actin filaments are locally redistributed and remodeled during AE (Figure S6B).

### Midpiece contraction in sperm located within the perivitelline space

In mice, only sperm undergoing AE prior to binding to the zona pellucida can penetrate and fertilize (*27*). We hypothesized that midpiece changes and motility cessation occur only after acrosome-reacted sperm penetrate the zona pellucida. Live imaging was performed after *in vitro* fertilization (IVF) using transgenic EGFP-DsRed2 sperm loaded with FM4-64. Eggs were inseminated with a high sperm count (200,000 cells) to increase the number of cells observed.

Figure 6A and Supplementary movie S7 show a sperm swimming within the perivitelline space. The FM4-64 fluorescence of this sperm midpiece is low (Figure 6B) which coincides with the fact that the sperm is moving (see also Figure 1F-G). Another example of this observation is shown in Figure 6C-D, an acrosome-reacted moving sperm within the perivitelline space had low FM4-64 fluorescence in the midpiece (Figure 6C). After 20 minutes, these sperm stopped moving and exhibited increased FM4-64 fluorescence, indicating midpiece contraction (Figure 6D). These results suggest that midpiece contraction and motility cessation occur after acrosome-reacted sperm penetrate the zona pellucida.

**Figure 6.**
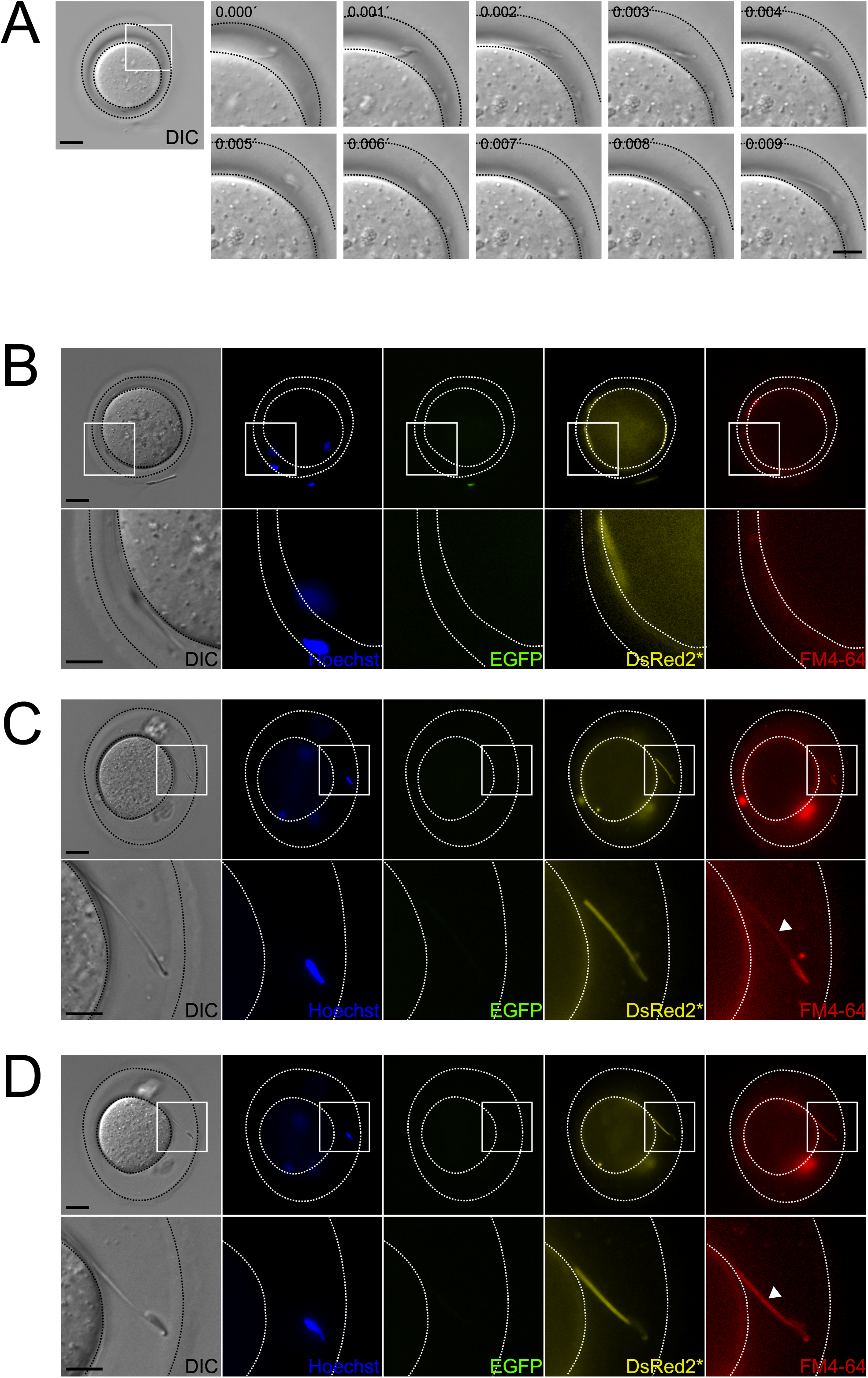
Occurrence of midpiece contraction in sperm located within the perivitelline space. Representative images of IVF experiments using EGFP-DsRed2 sperm. Oocyte-sperm complexes were stained with 10 μg/ml Hoechst and 10 μM FM4-64. A) Representative time series of DIC images showing a sperm moving within the perivitelline space. Scale bar in right panel = 20 μm, scale bar in left panel = 10 μm. B) DIC, Hoechst, EGFP, DsRed2*, and FM4-64 images are shown for the case depicted in Figure 6A, note that, as the sperm is moving, it is located in a different position in the perivitelline space. The area depicted in the upper panel is shown in higher magnification in the lower panel. Scale bar in upper panel = 20 μm, scale bar in lower panel = 10 μm. C-D) DIC, Hoechst, EGFP, DsRed2*, and FM4-64 images are shown for a (C) sperm that have passed through the ZP, displaying AE with an initially non-contracted midpiece. After 20 minutes, as shown in (D), the midpiece becomes contracted. The area depicted in the upper panel is shown in higher magnification in the lower panel. Scale bar in upper panel = 20 μm, scale bar in lower panel = 10 μm. Representative images from at least 6 independent experiments are shown. A total of 23 oocytes and 69 sperm were analyzed.

### Midpiece contraction takes place following sperm-egg fusion

Motile acrosome-reacted sperm, initially lacking midpiece contraction, can penetrate the zona pellucida. However, they eventually stop moving and exhibit an increase in FM4-64 fluorescence. This suggests that the cessation of motility plays a crucial role in fertilization events occurring after zona pellucida binding and penetration, and that a similar change in midpiece architecture is necessary for successful sperm-egg fusion. To explore this hypothesis, we designed a live imaging experiment using zona-free eggs. Denuded oocytes were loaded with Hoechst 33342, a nuclear dye, to visualize the exact moment of sperm-egg fusion (*28–30*). The experiment involved FM4-64, transgenic EGFP-DsRed2 sperm, and simultaneous signal collection from five separate channels (DIC, Hoechst, EGFP, DsRed2, and FM4-64). To enhance observation likelihood, volumetric data was collected by imaging at different z-planes (Figure 7A).

**Figure 7.**
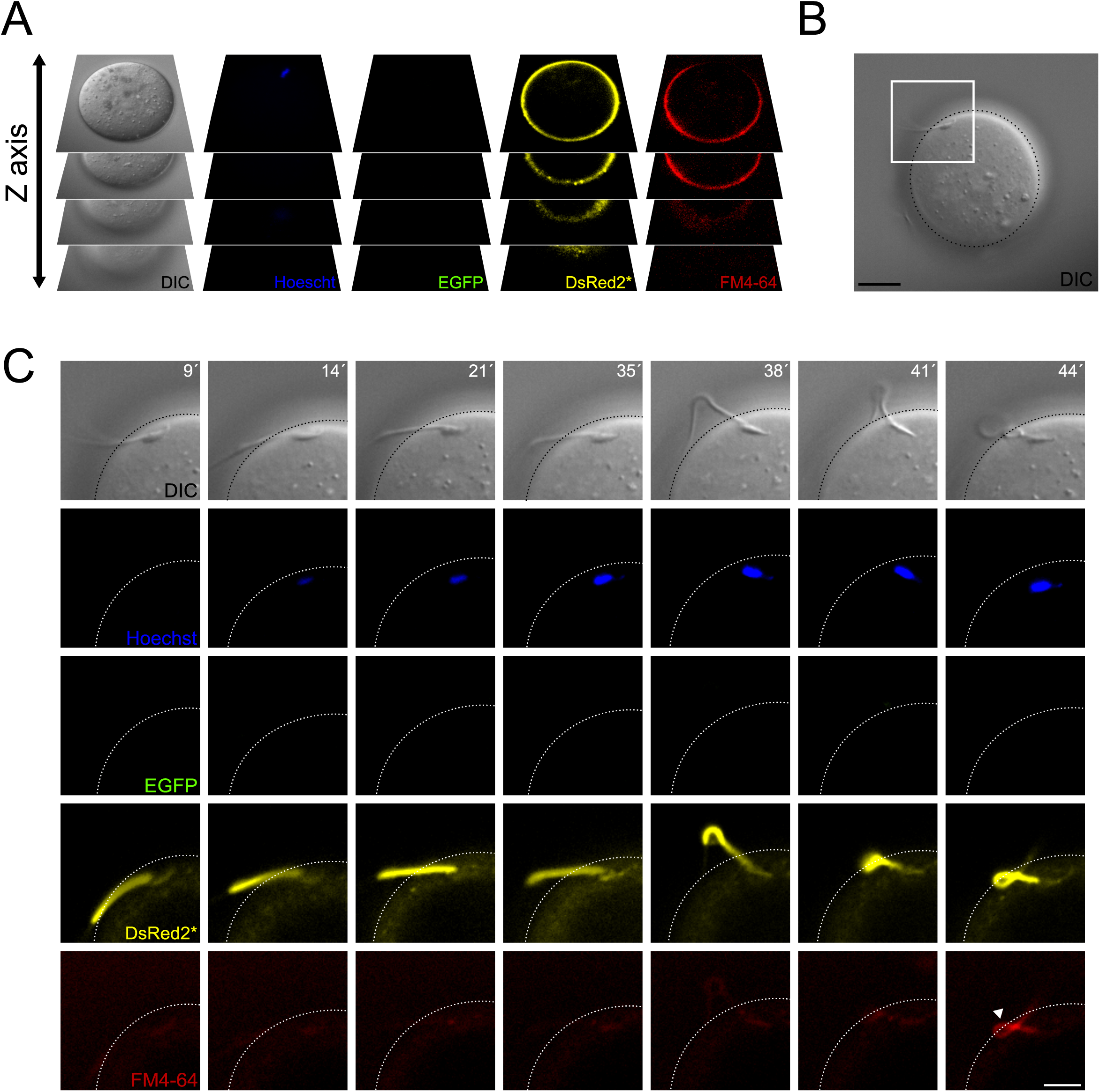
Contraction of the midpiece occurs after sperm-egg fusion. A) Schematic representation of the acquisition settings for the sperm-oocyte fusion assay. Images were taken every 7 μm along the z-axis. B) DIC image of a sperm-oocyte complex, with the area depicted in higher magnification in panel C. Scale bar = 20 μm. C) Representative time series of sperm-oocyte fusion assay experiments using EGFP-DsRed2 sperm. Oocytes were stained with 1 μg/ml Hoechst and 10 μM FM4-64. DIC, Hoechst, EGFP, DsRed2*, and FM4-64 images are shown over time. Scale bar = 10 μm. Note that midpiece contraction occurs after sperm-egg fusion and is proportional to the increase in FM4-64 fluorescence, as shown in Figure S2D, highlighting its potential importance in the fertilization process. Representative images from at least 4 independent experiments are displayed.

Figure 7C presents a representative volumetric time series of sperm-egg fusion, with only the optimal focal plane shown. The acrosome-reacted sperm (EGFP negative) initially did not display Hoechst fluorescence (Supplementary movie S8), indicating a lack of fusion. However, 14 minutes later, the dye entered the sperm and stained the nucleus, indicating fusion initiation. At this stage, sperm remain motile while bound to the egg plasma membrane. The midpiece diameter remained unchanged during initial sperm-egg fusion indicated by the low FM4-64 fluorescence. Following gamete fusion initiation, FM4-64 fluorescence increased (38 to 44 min), causing sperm motility to cease. In some instances, the midpiece folded and extended again (Figure S7B). These findings demonstrate that midpiece contraction occurs following sperm-egg fusion.

### A decrease in [Ca^2+^]_i_ in the midpiece following fusion precedes the midpiece contraction

In previous experiments, an increase in [Ca**^2+^**]_i_ was observed throughout the midpiece, coinciding with midpiece contraction (Figure 4 and Figure S3). To examine the mechanism behind midpiece contraction after fusion, wild-type sperm loaded with Fluo4 were exposed to denuded oocytes loaded with Hoechst. We hypothesized that dynamic changes in [Ca**^2+^**]_i_ during gamete fusion drive the midpiece contraction during sperm immobilization.

Figure 8A presents representative images from a time course live imaging experiment designed to observe sperm-egg fusion. The experiment involved simultaneous signal collection from four separate channels over time (DIC, Hoechst, Fluo4, and FM4-64). To enhance observation likelihood, images were taken at different z-planes, though only the optimal focal plane is shown.

**Figure 8.**
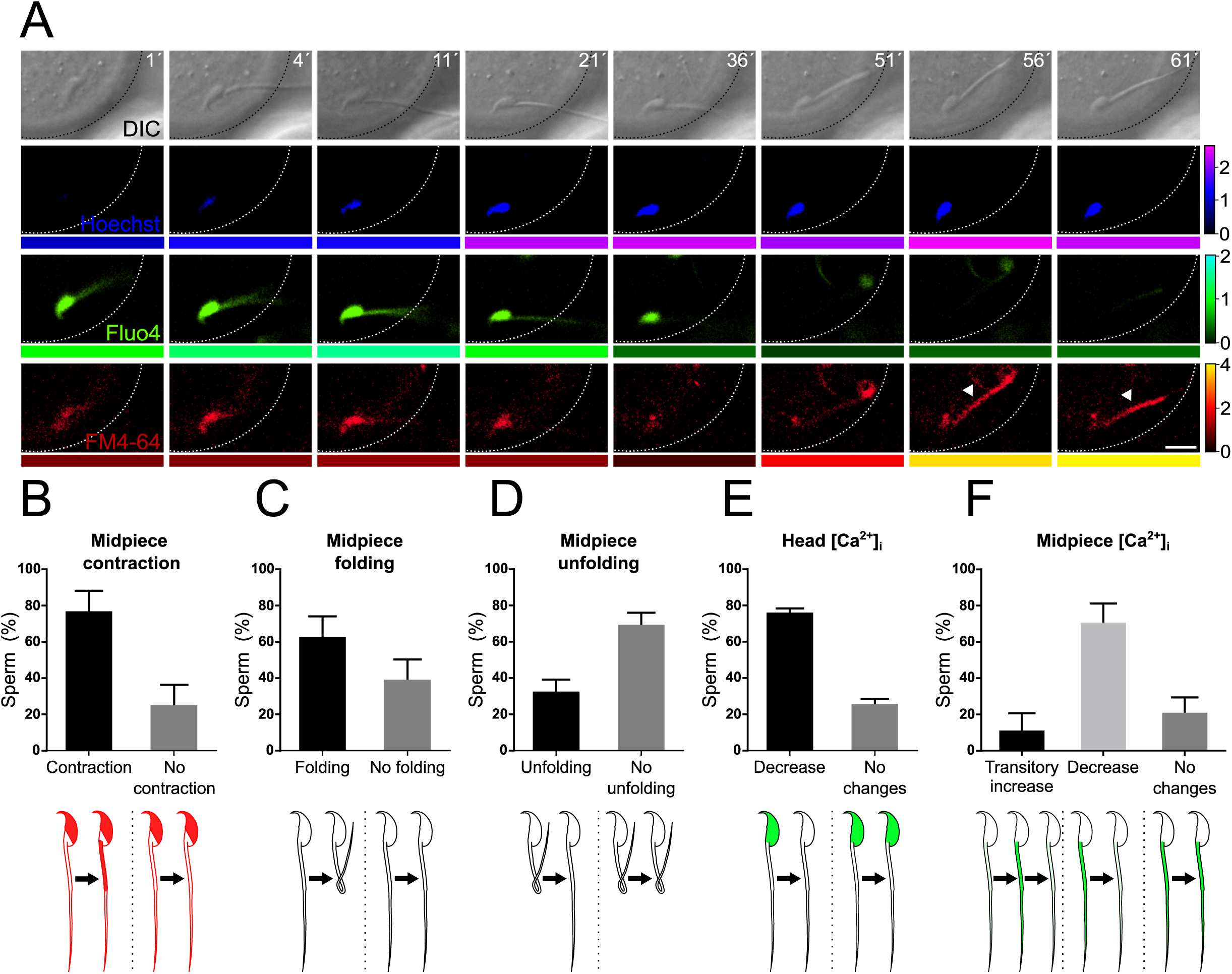
Midpiece contraction occurs in sperm-egg fusion after a decrease in [Ca^2+^]_i_. A) Representative time series of sperm-oocyte fusion assay experiments using wild-type sperm loaded with 1 μM Fluo-4. Oocytes were stained with 1 μg/ml Hoechst and 10 μM FM4-64. DIC, Hoechst, Fluo-4, and FM4-64 images are shown over time. Scale bar = 10 μm. The color code below each frame in the Hoechst (shown in blue), Fluo-4 (shown in green), and FM4-64 (shown in red) images indicates the normalized intensity of the fluorescence signal (scale bar on the right of the panel). B-F) Quantification of sperm showing midpiece contraction (B, indicated by increased FM4-64 fluorescence), midpiece folding (C), midpiece unfolding (D), Fluo-4 fluorescence dynamics in the head (E), and different patterns in the midpiece (F) during fusion. Data are presented as the mean ± SEM of the percentage of sperm counted for each experiment. Representative images and data from at least 3 independent experiments are shown. A total of 74 oocytes and 136 sperm were analyzed. Note that midpiece contraction occurs in sperm-egg fusion after a decrease in Fluo-4 fluorescence.

Unexpectedly, sperm binding to the egg plasma membrane displayed high levels of [Ca**^2+^**]_i_ in the head and midpiece (Figure 8A, time 1 min, and Supplementary movie S9). A few minutes after fusion began (indicated by an increase in Hoechst signal in the sperm), a decrease in [Ca**^2+^**]_i_ in the head and midpiece was observed (21 min to 51 min). This decrease in [Ca**^2+^**+]_i_ was followed by midpiece contraction, as evidenced by the increase in FM4-64 fluorescence (56 min). In several cells, the midpiece folded back before contracting. The gametes that underwent this change in the flagellum typically remained folded, but occasionally the tail unfolded again. Figures 8B-F quantify the key events described above during sperm-egg fusion.

In summary, 75.88 ± 12.20 % of the sperm exhibited midpiece contraction upon fusion. In 61.83 ± 12.04 % of cases, the midpiece folded on itself, and afterward, only 31.60 ± 7.47 % unfolded and stretched out again. All sperm bound to the plasma membrane before fusion presented high [Ca**^2+^**]i in the head, and the majority displayed a decrease in head [Ca**^2+^**]_i_ before midpiece contraction (75.13 ± 3.05 %). Concerning the midpiece, three patterns were observed during fusion: in the majority of cases (69.67 ± 11.46 %), the midpiece experienced a decrease in [Ca**^2+^**]_i_. Additionally, 10.33 ± 10.33 % of sperm showed a transient increase in [Ca**^2+^**]_i_, while the remaining 20.00 ± 9.41 % displayed no changes.

Collectively, these results indicate that a decrease in [Ca**^2+^**]_i_ in the midpiece after fusion precedes midpiece contraction and the cessation of sperm motility that precedes sperm-egg fusion.

## Discussion

In this article, we demonstrate the existence of a fundamental structural change that occurs in the sperm flagellum at the time of fusion with the egg. Using a plethora of advanced microscopy methods and single cell imaging, we provide insight about cellular and molecular events that occur in acrosome-reacted sperm, which, undoubtedly, are the ones capable of fertilizing an oocyte (*2*, *3*). At this precise moment of the fertilization process, sperm need to stop moving to complete the fusion between the two gametes. The cease of movement is caused by two concomitant processes that take place in the flagellar midpiece region: a change in the F-actin helical structure and a decrease in the midpiece diameter. To the best of our knowledge, this is the first time that a structural modification of the sperm flagellum related to a specific cellular necessity, i.e., gamete fusion, is described.

To arrive at the site of fertilization within the female reproductive tract, sperm motility is required. It is also fundamental to penetrate the different layers surrounding the egg. In this journey, sperm sense the environment and adapt their movement to the different physiological scenarios. In addition, recent evidence also demonstrated that after being released from their site of storage in the oviductal isthmus, mouse sperm undergo AE (*2*, *3*). This process occurs in the upper segments of the oviduct before any interaction with the eggs or their surrounding layers. This observation opened a new scenario about the motility of sperm in their last transit to the site of fertilization. Little is known about sperm in that region and how sperm move after AE. In this regard, most of the information about mammalian sperm comes from studies conducted *in vitro*, using a mixture of acrosome-intact and acrosome-reacted sperm. Even if their motility is analyzed in a subjective manner or using cell tracking systems, virtually all the experiments are conducted without discerning the acrosomal status of the cells. Another possible source of artifacts in this analysis is related with the fact that most of the experiments are also performed using epididymal sperm (not ejaculated) in aqueous solutions that support sperm capacitation but do not represent the natural viscous environment present in the female tract.

By performing in vitro experiments, we detected a strong association between the decrease in the midpiece diameter and the cease of sperm motility. This is observed in a subset of sperm after the occurrence of AE. This may suggest that only sperm that undergo AE in the oviduct and do not experience this midpiece contraction are capable of migrating to the ampulla and penetrate the cumulus and the zona pellucida. Those sperm that undergo this phenomenon earlier in the tract may not be suitable to continue their journey suggesting that this also may select the gametes during their transit to the ampulla. Previous observations tracking sperm within the female tract have shown the existence of acrosome-reacted sperm within the tract that remain non-motile (*2–4*).

Acrosome-reacted sperm that bind and penetrate the zona pellucida are ready to fuse with the egg. During sperm-egg fusion, several authors have reported in different species that sperm stop moving (*7–9*). However, the mechanism behind this behavior is not established. One possible explanation is that fusion promotes ion transport changes in sperm, which significantly alter flagellar movement. In this sense, it was previously demonstrated that certain ions such as Ca^2+^ may diffuse from the oocyte to the sperm (*31*). However, a clear technical limitation in our experimental approach is that the probes that monitor ion dynamics may be exchanged between both gametes. If the concentration of a given probe does not remain stable, it is impossible to determine the accurate change that occurs during fusion. Future experiments may take advantage of transgenic models that incorporate a particular sensor to study this process in vivo, such as the one used by Cohen and collaborators (*32*).

Regardless of the precise nature of ion exchange between sperm and eggs, the diameter of the sperm flagellum in the midpiece is reduced. This midpiece diameter reduction is strongly associated with cessation of sperm motility and is apparently needed to complete the fusion process. Our observation clearly supports this notion. Importantly, the sperm flagellum folds back during fusion coincident with the decrease in the midpiece diameter. Interestingly, this was previously observed in mammals as well as in sea urchin sperm (*7*, *33*, *34*).

Like other cylindrical biological structures, the sperm flagellum relies on the cytoskeleton for its structural organization and specialized mechanical properties. In addition to the change in midpiece diameter, a significant rearrangement of the F-actin cytoskeleton also occurs. In the midpiece, the polymerized actin is organized in a double helix accompanying the mitochondria (*13*). As in many other organisms, the actin cytoskeleton possesses important structural functions, and dynamic changes of F-actin allow the cells to conduct important cellular tasks such as exocytosis. However, less is known about the structural changes undergone by the actin cytoskeletons in cilia and flagella. Remarkably, it has been reported that actin forms helix-like structures in the flagellum of the parasite *Giardia intestinalis* (*35*). These findings open the question of whether, to some extent, flagellar helical structures are conserved among diverse species. Our observations demonstrate that the F-actin double helix undergoes a change in the helical pitch as well as in the radial distance to the axoneme. This is concomitant with the decrease in midpiece diameter. Our single-cell experiments using super-resolution microscopy also revealed that the plasma membrane approached the F-actin network during this change. It is well established that various proteins can function as linkers between the plasma membrane and the actin cytoskeleton (*36*), but their roles in this specific process remain to be studied. Regardless of how these structures are connected, it is evident that both are associated. However, our experimental data cannot determine whether the plasma membrane is causing the change of the actin network or if the actin network influences the plasma membrane.

Another observation emerging from our study is that a change in [Ca**^2+^**]_i_ occurs prior to the midpiece contraction. It is well known that Ca**^2+^** is important for AE, and a specific transient rise in [Ca**^2+^**]_i_ originating in the head can trigger exocytosis (*19*). Previous observations have demonstrated that the sperm head and tail are not isolated compartments, and that ions and other molecules can move between them (*37*, *38*). In our single-cell experiments, we observed that the rise in FM4-64 fluorescence in the midpiece occurs after the increase in [Ca**^2+^**]_i_ in that region, suggesting a potential association between these events. This could involve diffusion or active transport processes; further investigation is required to determine the precise mechanism to demonstrate if the structural changes are triggered by Ca2+. In addition, [Ca^2+^]_i_ increase and/or modification in the midpiece architecture may result in functional changes in the mitochondria such as the status of the mitochondrial membrane potential and the ATP production. This possibility needs to be further explored.

The same phenomenon was studied in sperm bound to the egg plasma membrane to evaluate if the rise in [Ca^2+^]_i_ also occurs at the time of fusion. Remarkably, we noticed that most of the bound sperm that ended up fusing with the eggs displayed high levels of [Ca^2+^]_i_ in both the head and the flagellum. The midpiece contraction and the immobilization occurred when the levels of [Ca^2+^]_i_ went down in the midpiece suggesting a possible connection between both events. However, our experimental approach limits the interpretation of this result. One possible explanation is that as soon as sperm bind to the plasma membrane of the oocyte, there is a rapid increase in [Ca^2+^]_i_ in the sperm. On the other hand, a massive transport of Ca^2+^ from the egg to the sperm could also occur. These hypotheses, however, are hindered by the technical limitations mentioned above. As the oocyte is not loaded with Fluo4, we cannot rule out that the apparent [Ca^2+^]_i_ decrease seen in these experiments is due to dye diffusion into the oocyte. Either way, it is clear that the transport of this ion into or out of the sperm is key to cease motility at this fundamental step of fertilization (*16*). Additionally, these results demonstrate that only sperm with elevated [Ca^2+^]_i_ are capable of binding to the eggs. All these possible scenarios need to be determined in future experiments. Thus, regardless of the mechanism, a clear change in [Ca^2+^]_i_ dynamics is observed at the time of midpiece contraction. Future studies will aid to indicate if certain Ca^2+^-dependent proteins that can modify the actin cytoskeleton are responsible for this change.

Why sperm stop moving: We propose three possible hypotheses. The cessation of sperm motility can be attributed to the simultaneous or not occurrence of various events. 1) a rapid increase in [Ca^2+^]_i_ levels may trigger the activation of Ca^2+^ pumps within the flagellum. This process consumes local ATP levels, disrupting glycolysis in the process. 2) Reorganization of the actin cytoskeleton: alterations in the actin cytoskeleton can lead to changes in the mechanical properties of the flagellum, impacting its ability to move effectively. 3) Midpiece contraction: Contraction in the midpiece region can potentially interfere with mitochondrial function, thereby impeding the energy production necessary for sustained motility. In addition, we speculate that the folding of the flagellum during fusion further facilitates sperm immobilization because it makes it more difficult for the flagellum to beat. Such process can enhance stability and increase the probability of fusion success. Mechanistically, the folding may occur as a consequence of the deformation-induced stress that develops during the decrease of midpiece diameter.

In cilia, apart from the findings presented in this paper, nothing is known about the regulation of motility by actin. Actin has been found to participate in ciliogenesis, but its involvement in active motility regulation has not been reported. This highlights a potentially unique role for the actin cytoskeleton in regulating sperm function during fertilization. Our discovery of actin’s dynamic reorganization in sperm suggests it could have a more active role in regulating motility and other functions. Given the conservation of cilia and flagella across various organisms, our discovery could prompt further research on the role of the actin network in these structures.

In conclusion, we demonstrate that sperm undergo a structural reorganization of the actin cytoskeleton during key events of fertilization, as summarized in the working model shown in Figure 9. Our findings introduce a new aspect of study in reproductive biology. Previous research has mainly focused on identifying proteins essential for sperm-egg fusion. Our results reveal a previously unexplored biological mechanism in mammalian fertilization, opening new avenues for contraceptive method development.

**Figure 9:**
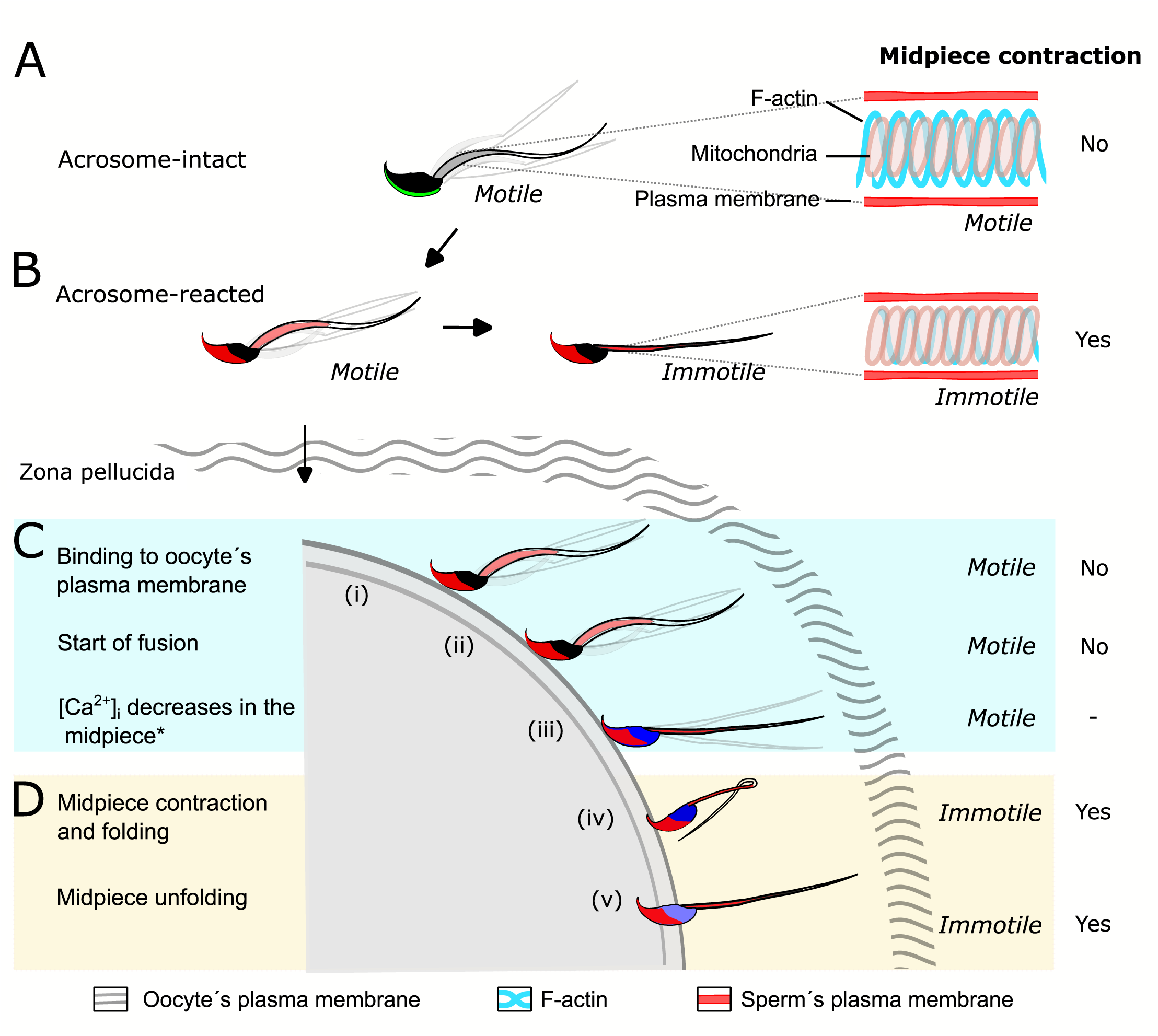
Proposed model of the structural reorganization of the sperm actin cytoskeleton during key events of fertilization. The double helix actin network surrounding the mitochondrial sheath of the midpiece undergoes structural changes prior to the motility cessation. This structural modification is accompanied by a decrease in diameter of the midpiece and is driven by intracellular calcium changes that occur concomitant with a reorganization of the actin helicoidal cortex. Although midpiece contraction may occur in a subset of cells that undergo AE (A and B), the midpiece contraction occurs prior to motility cessation observed after sperm-egg fusion (C and D).

## Materials and methods

Detailed methods and protocols are provided in the SI appendix. For sample details, optical equipment, imaging conditions and probes used, see Supplementary Table S1.

## Acknowledgments

We gratefully acknowledge the financial support provided by the Williams and Rene Baron Foundations, the Male Contraceptive Initiative (MCI), and the Chan-Zuckerberg Initiative (CZI) for this work. Our sincere thanks go to Dr. Pablo Visconti for his valuable insights throughout the course of this project. We also thank Yoloxochitl Sánchez-Guevara for her technical support. This paper is dedicated to the memory of OAP.

## Author contributions

Investigation: MJ, GML, MDGE, CSC, JLDLVB, AL, AAH, MPRG, ALS. Data curation: MJ and VXAA. Writing: MJ, MGB and AG. Methodology: XX, Diego K and GK. Conceptualization: Dario K, Diego K, AD, MGB and AG. Supervision: MGB and AG.

## Data, Materials, and Software Availability

Data acquired and analyzed during this study are included in this manuscript or will be available at Zenodo. Github will be used for depositing the code use in analysis.

