## Supplementary figures and images for "Reorganization of the Flagellum Scaffolding Induces a Sperm Standstill During Fertilization"

### Supplementary Figures

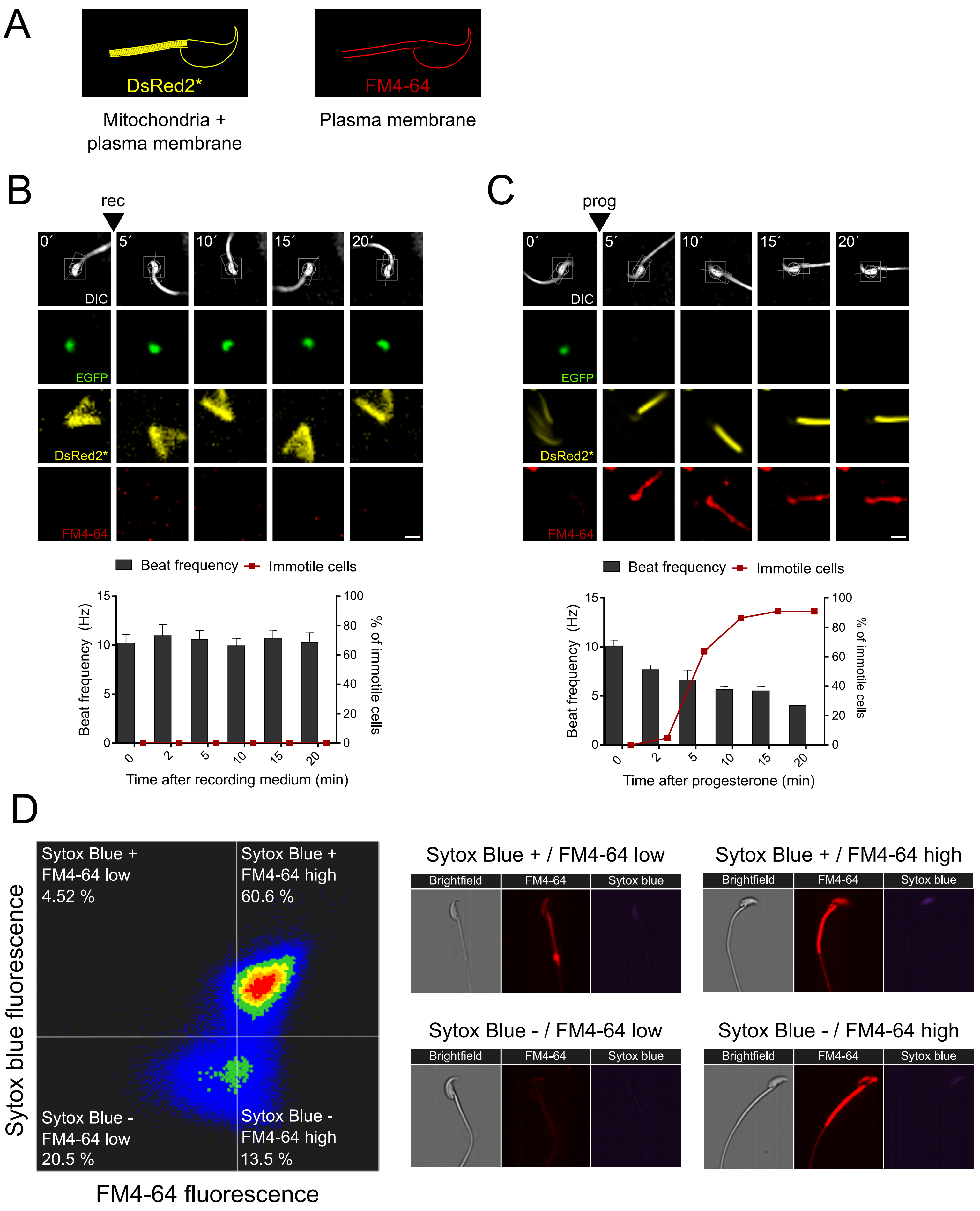

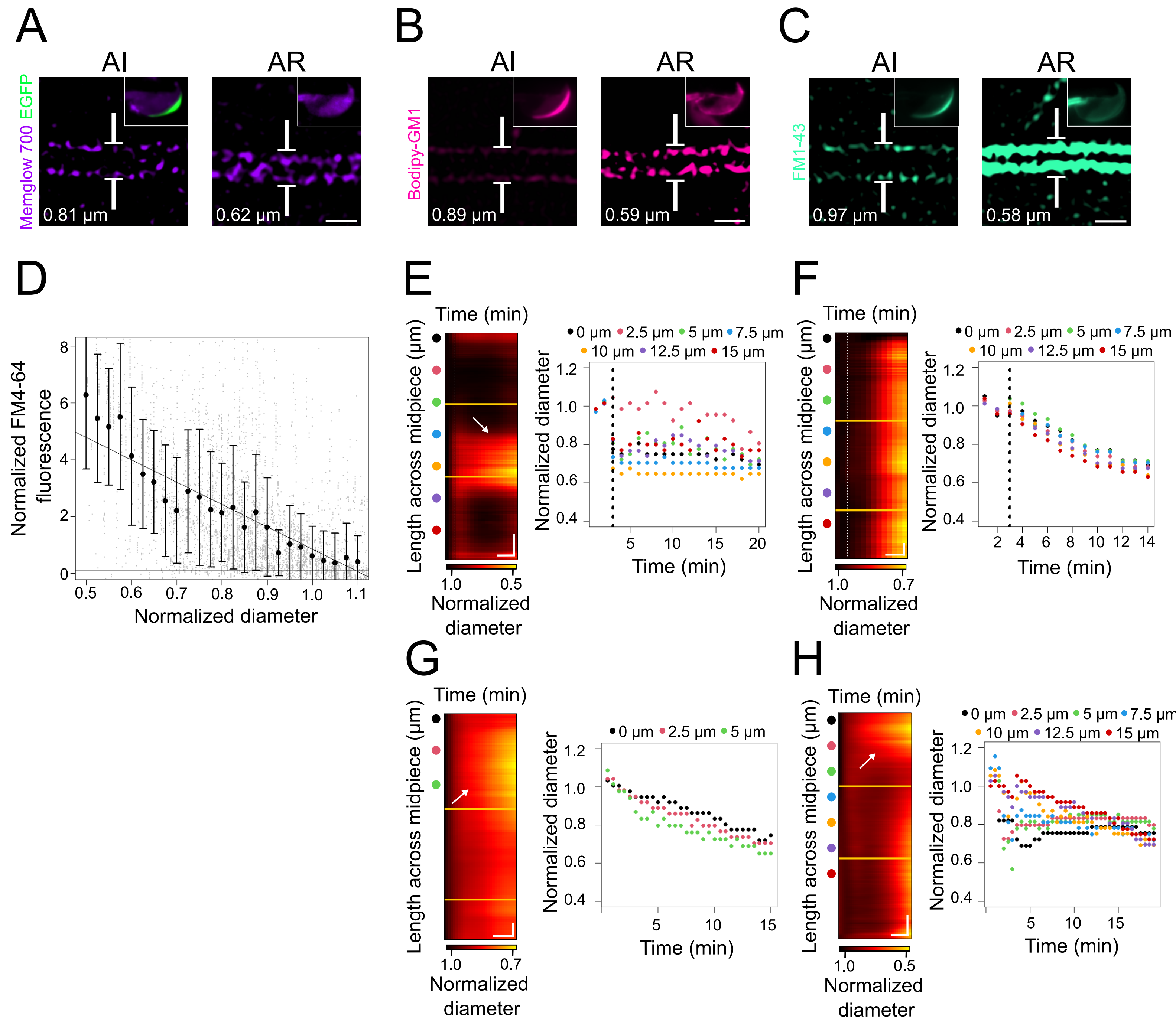

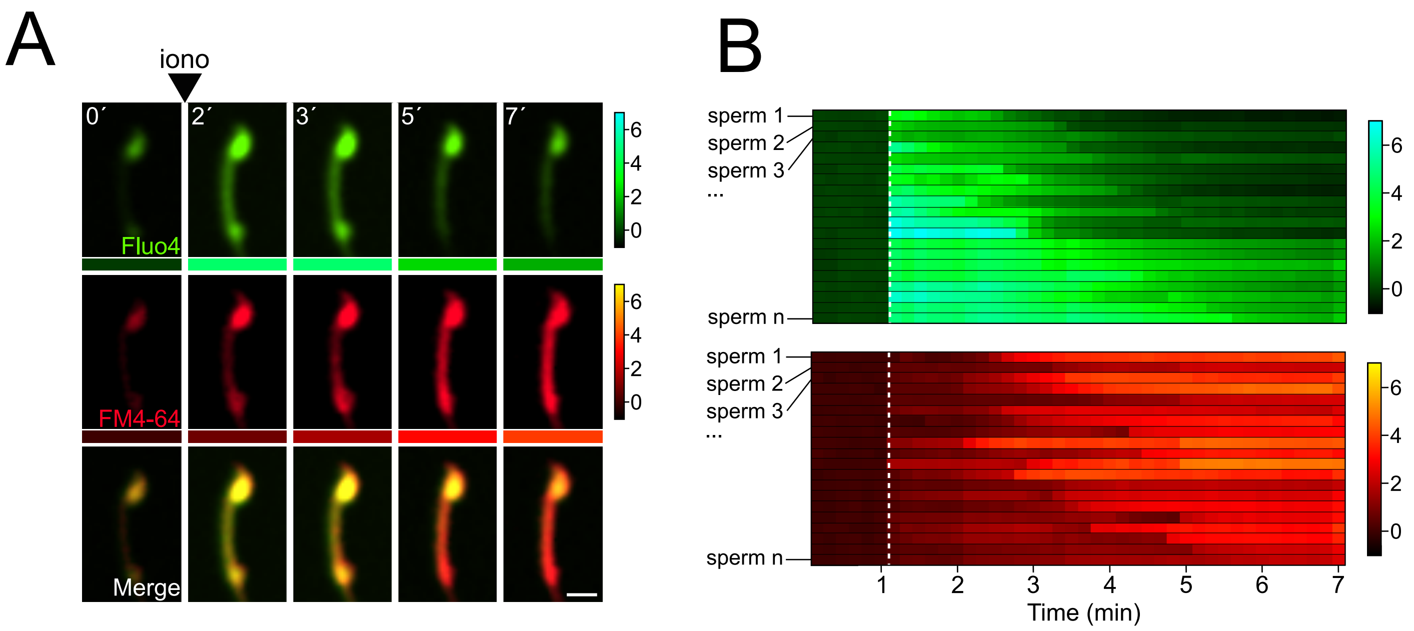

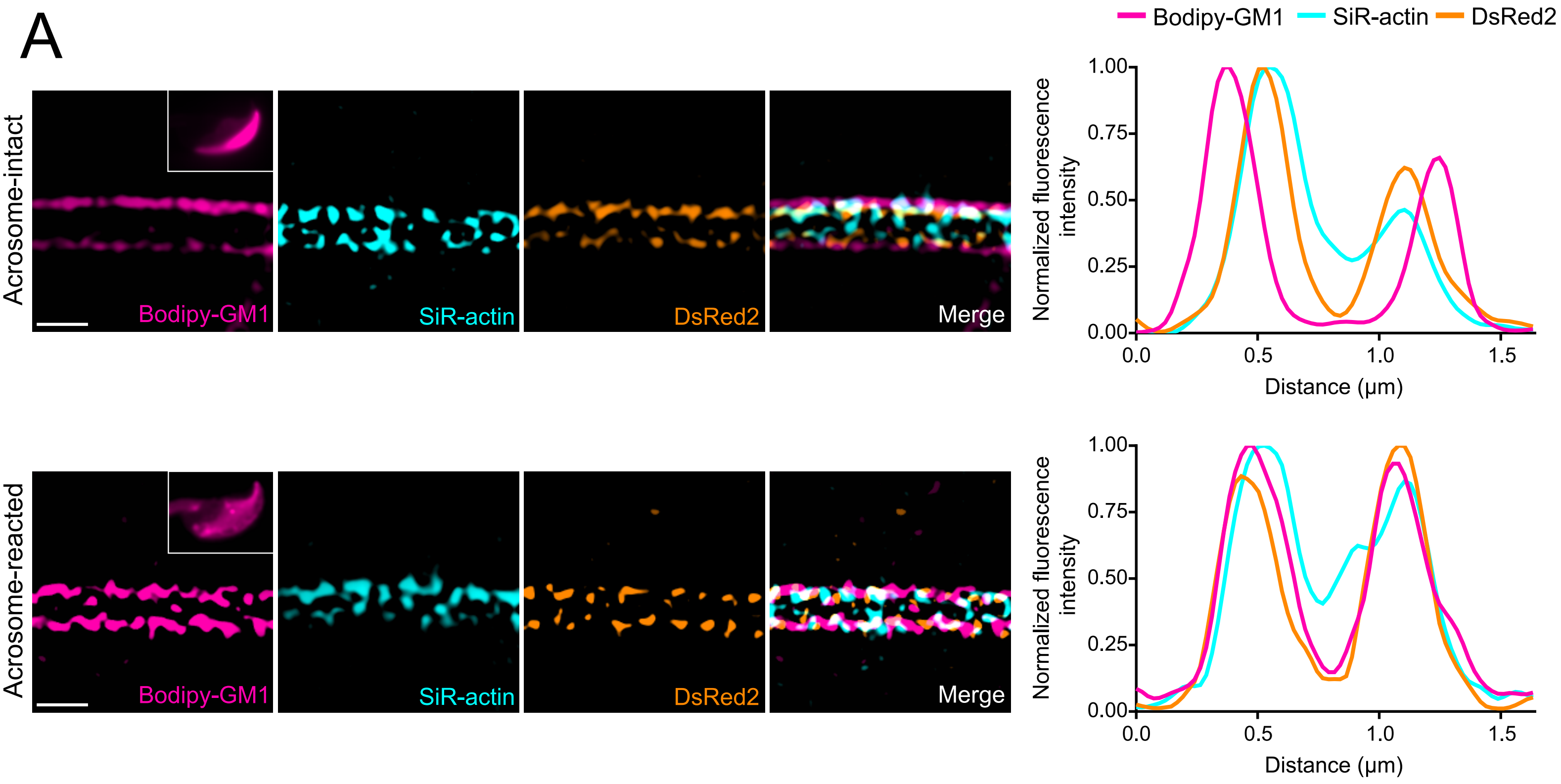

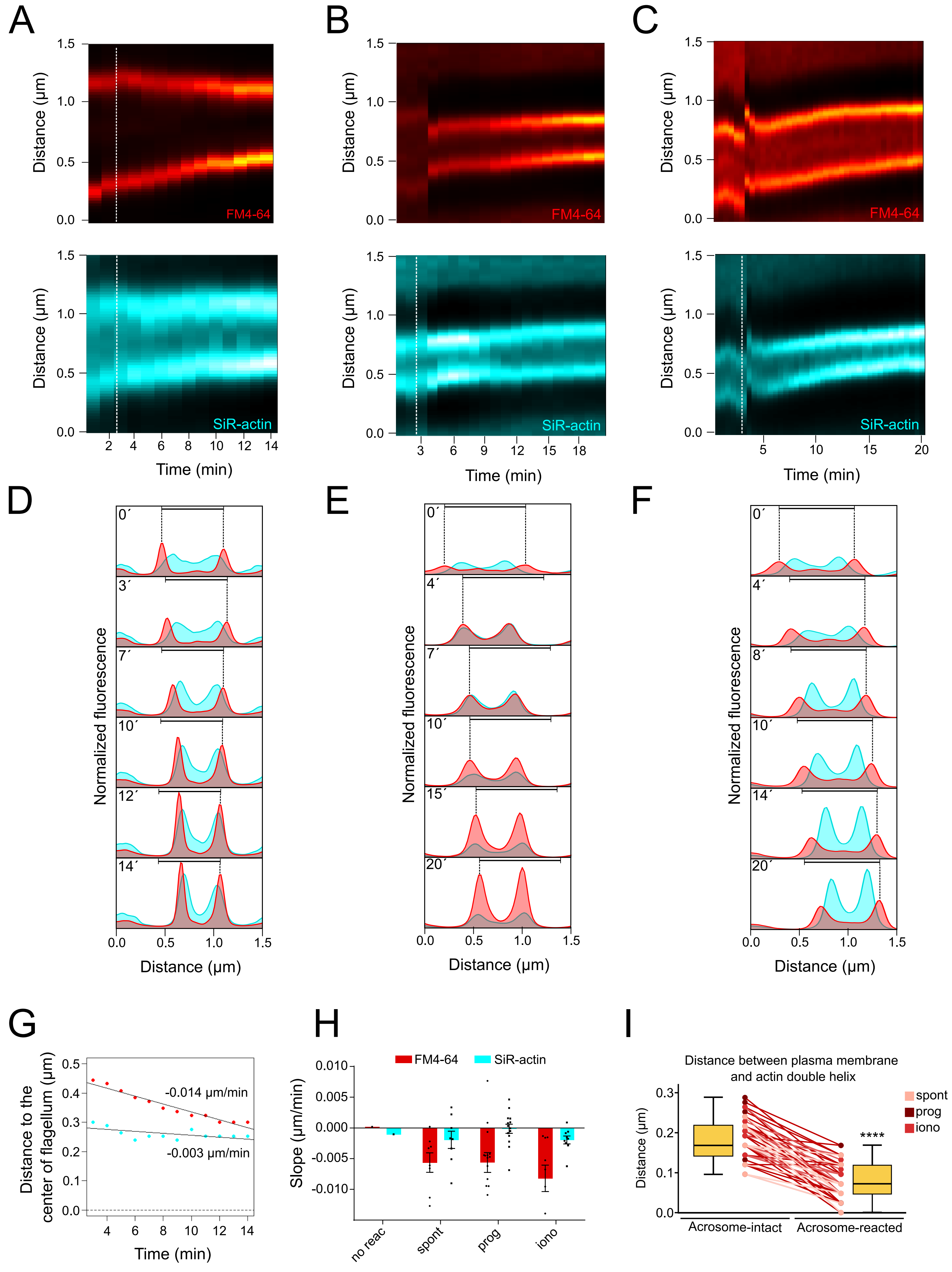

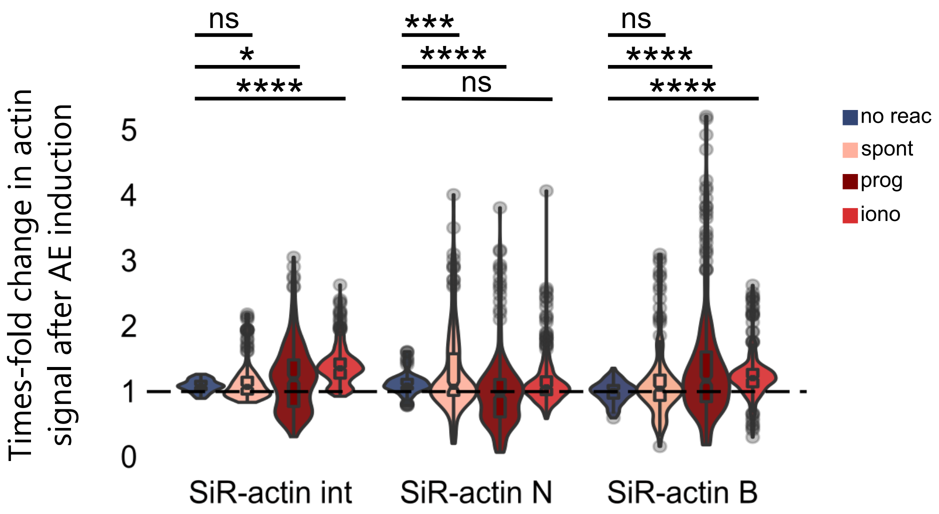

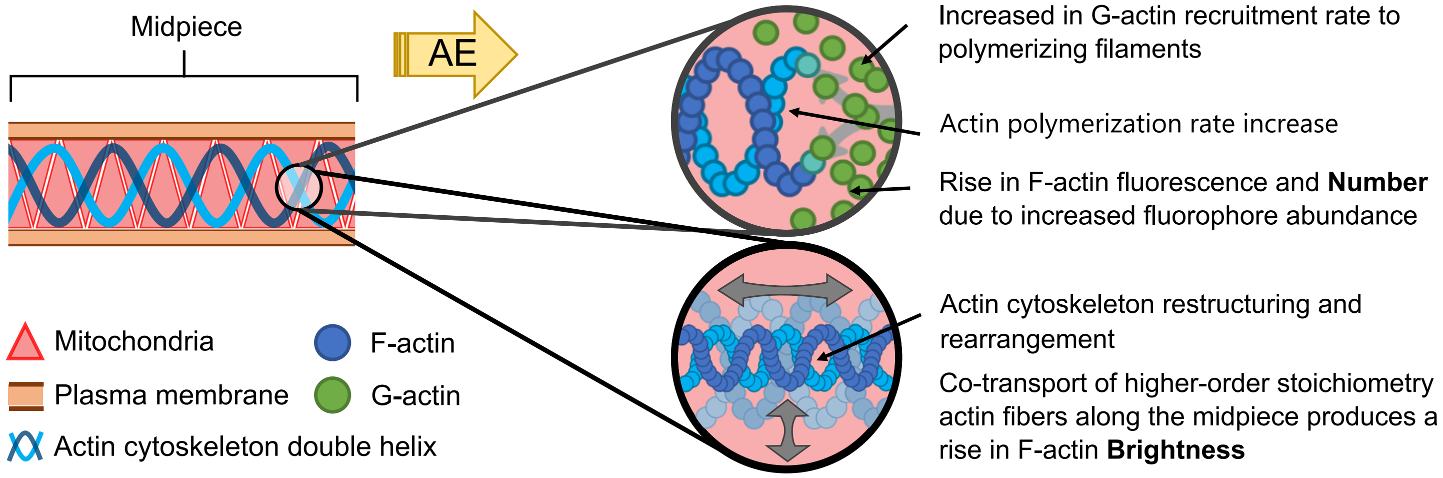

#### Figure S7

A

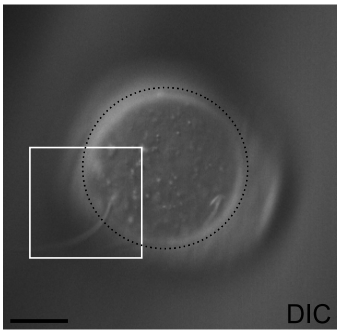


B

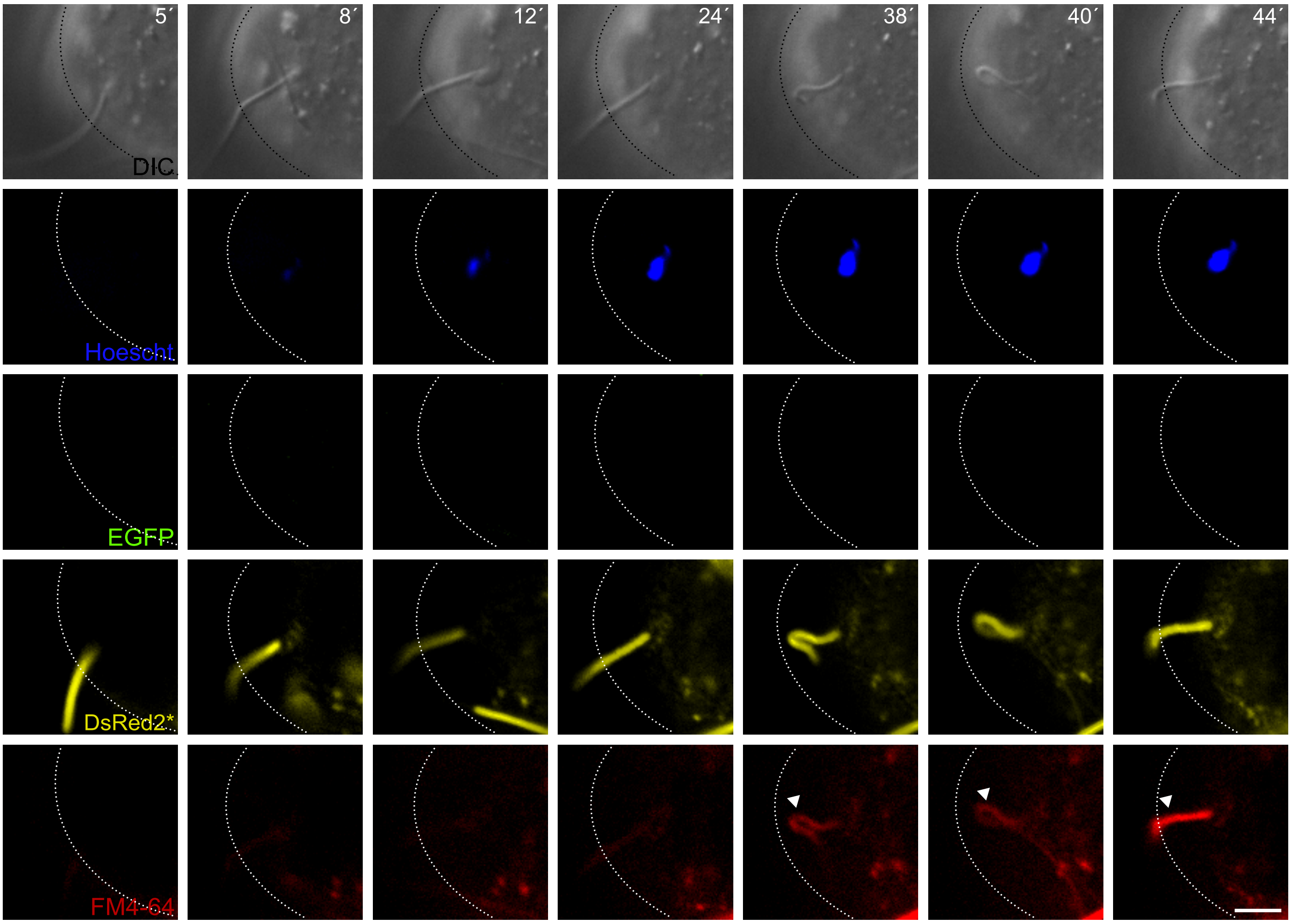

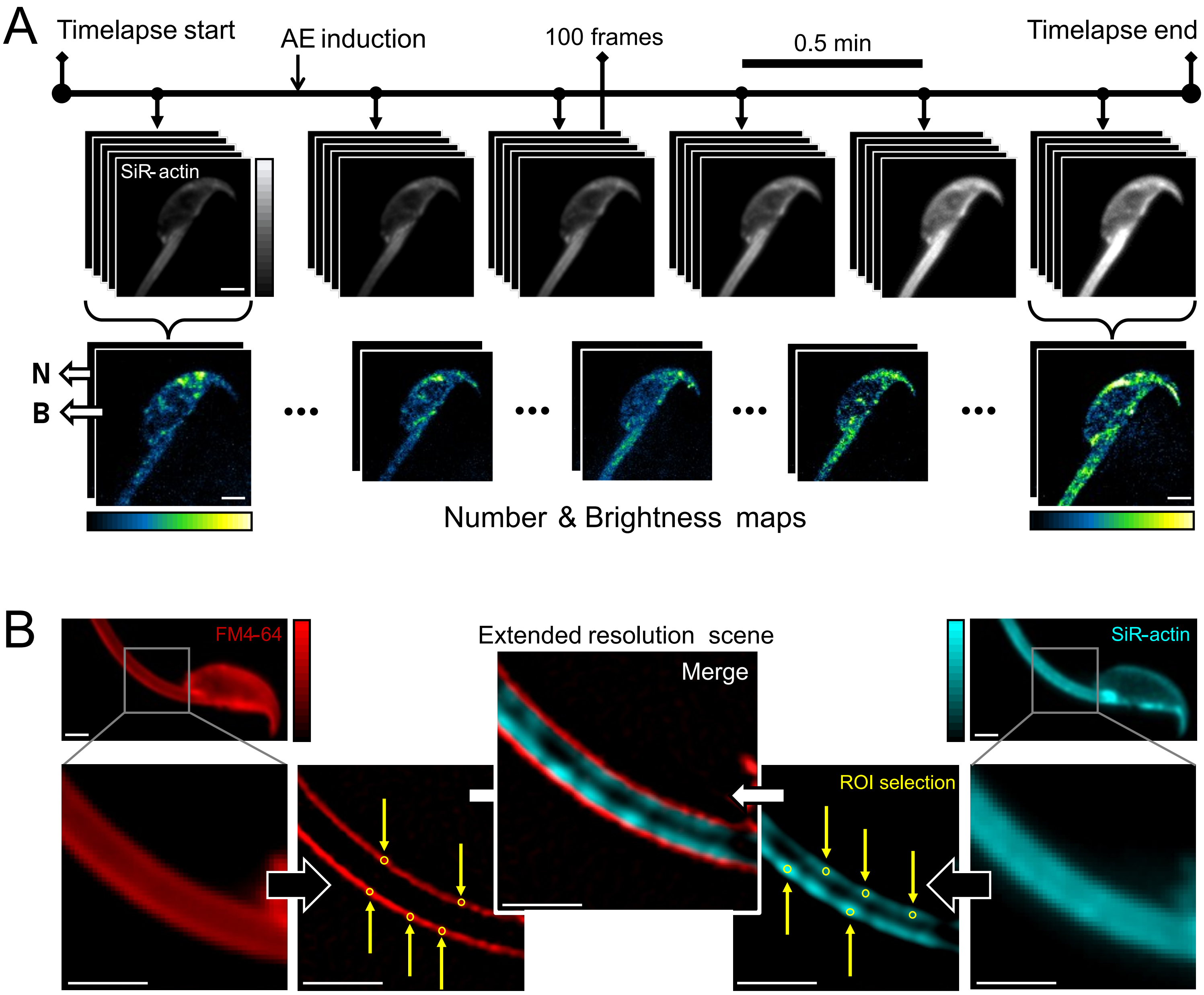
