## Supplementary material for "Reorganization of the Flagellum Scaffolding Induces a Sperm Standstill During Fertilization": Materials and methods

**Reagents and chemical sources**

The chemicals used in this study were procured from the following sources: Progesterone, laminin, and concanavalin A were obtained from Sigma-Aldrich Chemical Co. (St. Louis, MO, USA). Fluo-4 AM, pluronic acid, Alexa Fluor-647–phalloidin, FM4-64, FM1-43, Hoechst 33342, and Bodipy-GM1 were acquired from Invitrogen, Thermo Fisher Scientific (Waltham, MA, USA). SiR-Actin and Memglow 700 were sourced from Cytoskeleton (Denver, CO, USA), while ionomycin was purchased from Cayman Chemicals (Ann Arbor, MI, USA). All other chemicals utilized in this research were of reagent grade.de.

**Animals and housing conditions**

Mature mice (8- to 12-week-old) from the following strains were used in this study: CD1, Hybrid F1 (Balb/C x C57BL/6), and transgenic B6D2F1-Tg (CAG/mt-DsRed2, Acr-EGFP) RBGS002Osb (*39*). Mice were housed in groups of four in a temperature-controlled environment maintained at 23°C, with a light cycle from 07:00 to 19:00 h. The animals had *ad libitum* access to tap water and laboratory chow.

All experimental procedures adhered to the guidelines of the Institutional Animal Care and were reviewed and approved by the Ethical Committee of the Instituto de Biotecnología, UNAM, and the Instituto de Biología y Medicina Experimental, Buenos Aires. The experiments were conducted in strict accordance with the Guide for Care and Use of Laboratory Animals approved by the National Institutes of Health (NIH).

**Sperm medium and capacitation**

The non-capacitating medium employed in this study was a modified TYH medium, which consisted of 119.3 mM NaCl, 4.7 mM KCl, 1.71 mM CaCl_2_·2H_2_O, 1.2 mM KH_2_PO^4^, 1.2 mM MgSO_4_·7H_2_O, 0.51 mM sodium pyruvate, 5.56 mM glucose, 20 mM 4-(2-hydroxyethyl) piperazine-1-ethanesulfonic acid (HEPES), and 10 μg/ml gentamicin (NC medium). To create capacitating conditions, 15 mM NaHCO_3_ and 5 mg/ml BSA were added to the medium (CAP medium). In all experiments, the pH was adjusted to 7.4 using NaOH

The animals were euthanized, and cauda epididymal sperm were harvested. Both cauda epididymides were placed in 1 ml of NC medium. After a 15-minute incubation at 37°C (swim-out), the epididymides were removed, and the sperm were resuspended to a maximum final concentration of 10^7^ cells/ml in 100 μl of the appropriate medium. Subsequently, an equal volume (100 μl) of either non-capacitating medium or two-fold-concentrated capacitating medium (30 mM NaHCO_3_ and 10 mg/ml BSA) was added, and the sperm were incubated for 60 minutes at 37°C.

**Live imaging of transgenic mice sperm undergoing AE**

Once capacitated, sperm from transgenic B6D2F1-Tg mice were immobilized on coverslips coated with either concanavalin A or laminin (1 mg/ml) to facilitate recordings. Unattached spermatozoa were gently washed away, and the chamber was filled with the recording medium (NC medium) containing 10 μM FM4-64. The application of this dye for continuous AE recording and simultaneous measurement of other parameters, such as [Ca^2+^]_i_ concentration, has been previously established (*16*). Progesterone (with a stock concentration of 50 mM dissolved in dimethyl sulfoxide, DMSO), ionomycin (with a stock concentration of 2 mM dissolved in DMSO), or vehicle (DMSO) was prepared in the recording medium and gently applied using a micropipette.

**Sperm motility analysis**

For imaging, capacitated sperm from transgenic B6D2F1-Tg mice were immobilized on laminin-coated coverslips (1 mg/ml). Recordings were performed as described earlier, using a recording medium (NC medium) containing 10 μM FM4-64. The beating frequency of sperm flagella was calculated using the method proposed by Corkidi and colleagues (*40*). The flagella of individual sperm cells were tracked from recorded videos (2-minute duration at 100 frames per second) for each experimental condition. The mean orientation of the flagella was detected to infer their beating frequency through the Fourier transform. Each condition was assessed before and after the addition of progesterone, over a total duration of 20 minutes (with 2-minute videos captured every 5 minutes).

**Scanning electron microscopy (SEM)**

For the fixation of mouse spermatozoa for scanning electron microscopy (SEM), we utilized the following reagents and media: 0.1 M sodium cacodylate buffer, pH 7.4; TAGA mix, which contains 3.5% glutaraldehyde and 0.5% tannic acid. Coverslips were cleaned by immersing them in 70% ethanol, followed by a mixture of 1 M hydrochloric acid and 50% ethanol for 1 hour, and rinsed with Milli-Q water. Subsequently, they were dried and subjected to plasma cleaning in a Plasma Etch Inc. PE-25 Ver 1002 oven under specific conditions: vacuum set point of 150 mTorr, atmospheric vent for 45 seconds, purge vent for 5 seconds, gas stabilization for 15 seconds, and a vacuum alarm set to 3 minutes. Spermatozoa were obtained by dissecting the cauda epididymis from mice and resuspending them in 1 mL of NC TYH medium. The samples were incubated at 37°C for 15 minutes, after which 400 μL of medium was collected into two separate tubes. One tube was adjusted to a final volume of 800 μL with 2X CAP medium to achieve 1X CAP medium and incubated for 60 minutes, followed by the induction of the acrosome reaction using 30 μL of Ionomycin for 30 minutes. Both tubes were centrifuged at 300 G for 3 minutes, and the supernatant was carefully removed, leaving 100 μL. For fixation, the samples were diluted with 100 μL of PBS and 600 μL of 0.1 M sodium cacodylate buffer and placed on coverslips. After a 10-minute incubation, the coverslips were washed three times with 0.1 M sodium cacodylate buffer, each involving shaking in a digital rotator at 80 RPM for 10 minutes. Subsequently, 100 μL of TAGA mix was added to each coverslip for 15 minutes. Finally, the coverslips were preserved by submerging them in 0.1 M sodium cacodylate buffer within a petri dish until further SEM analysis using an Auriga-FIB-Zeiss scanning electron microscope.

To facilitate the quantification of the midpiece diameter of the mouse sperm flagellum, we manually annotated 157 well-focused Scanning Electron Microscopy (SEM) images. These annotations were used to create a comprehensive image dataset specifically for measuring the midpiece. We employed the ImageJ software for the annotation process. Using the “Polygon Selection Tool,” we carefully delineated the midpiece of each flagellum. Each image was consistently aligned so that the sperm head was positioned to the left, ensuring the midpiece was as straight as possible for accurate measurement. A polygon was drawn around the edges of the midpiece to define a Region of Interest (ROI). These ROIs were then saved in a compressed ZIP format. Subsequently, these saved ROIs were imported into a Python script executed in a Jupyter Notebook. This script enabled us to compute the average diameter of the midpiece across all images in the dataset. By leveraging Python’s image processing libraries, we automated the calculation, ensuring precision and reproducibility in our measurements.

**Imaging Flow Cytometry**

For the analysis of mouse spermatozoa under non-capacitating conditions using image cytometry, the viability dye, Sytox Blue, was procured from Invitrogen, Thermo Fisher Scientific (Waltham, MA, USA). Transgenic B6D2F1-Tg (CAG/mt-DsRed2, Acr-EGFP) RBGS002Osb mice were used. Sperm were retrieved by sacrificing mice and dissecting the cauda epididymides, which were incubated in 1 mL of NC TYH medium at 37°C for 15 minutes (swim-out). After incubation, approximately 800 μL of medium containing the sperm was collected, centrifuged at 3000 RPM for 3 min, and the volume was reduced to 300 μL to concentrate the cells. The samples were kept on ice to preserve viability. For fluorescence labeling, Sytox Blue stock solution (1 mM in DMSO) was diluted to 50 μM in TYH medium, and further adjusted to 1 μM with sperm samples using a final volume of 60 μL, mixing 58.8 μL of sperm suspension with 1.2 μL of Sytox Blue solution. Image acquisition was performed using an AMNIS imaging system with a 60x objective (NA 0.9) and lasers set at 120 mW (488 nm) and 140 mW (561 nm). Sperm images were captured for 30 minutes, focusing on cells using Aspect Ratio vs. Area in brightfield and RMS Gradient for in-focus selection. Segmentation and analysis of populations of interest were conducted in IDEAS software, where masks for brightfield and fluorescence images were applied to isolate live and dead populations based on Sytox Blue and acrosome fluorescence intensity (EGFP). The resulting data were exported as 16-bit raw images for further analysis.

**Single cell live imaging of midpiece membrane during AE**

Capacitated CD1 mouse sperm were recorded using the previously described method. The recording medium (NC medium) containing 0.5 μM FM4-64 was utilized for this process. The midpiece diameter and fluorescence intensity were quantified using ImageJ software version 1.47V (National Institute of Health, USA). For fluorescence intensity measurement, regions of interest (ROIs) were designated in the sperm head and midpiece. After subtracting the background, the formula (F-F_0_)/F_0_ was applied, where F_0_ represents the baseline calculated by averaging the frames prior to stimulus application.

**Analysis of diameter and FM4-64 fluorescence intensity using kymographs**

Kymographs were constructed with the ImageJ Kymograph plug-in. For super-resolution kymographs, a cross line was plotted every 2.5 μm along the midpiece in Super-Resolution Radial Fluctuations (SRRF) images. The R language environment (R 3.3.3 GUI 1.69 Mavericks build, 7328) was employed to obtain diameter values. The super-resolution kymographs were processed using the autocorrelation (acf) function to determine the positions of fluorescence maxima on both sides of the midpiece membrane. The distance between the maxima represented the diameter value. These values were then normalized to those obtained prior to the addition of AE stimulants or vehicle. For fluorescence kymographs, a line was drawn along the midpiece in Total Internal Reflection (TIRF) images. The fluorescence values were also acquired using the R language environment. After subtracting the background, the formula (F-F_0_)/F_0_ was applied, where F_0_ denotes the baseline, calculated by averaging the frames before stimulus application.

**Correlation between FM4-64 midpiece fluorescence and normalized diameter. Data transformation from fluorescence to contraction.**

The kymograph analysis provided two main values for each time point evaluated: normalized diameter values every 2.5 μm from the super-resolution kymographs, and normalized fluorescence values for each pixel analyzed (one value every 0.117 µm) from the fluorescence kymographs. To examine the relationship between these values, a graph was created in the R language environment that included all data points, encompassing all analyzed cells, all diameter measurements (every 2.5 μm), and their corresponding fluorescence values for all time points evaluated. In this graph, the x-axis represents the normalized diameter, while the y-axis represents the normalized fluorescence (Figure S2D). Additionally, a linear regression was performed using the general formula 𝑚𝑥+𝑏, where 𝑚 is the slope with a value of 8.90 ± 0.15, 𝑏 is the y-intercept with a value of -8.05 ± 0.17, and R2 is 0.32. With the parameters of the linear regression, we could approximate the midpiece diameter value based on a given FM4-64 fluorescence value. Despite data dispersion, this method provided an approximate value that could indicate whether the midpiece was contracting or not. It is noteworthy that the observed data dispersion may have biological significance at the cell population level, where heterogeneity among individual spermatozoa may contribute to variability in their physiological responses.

**Live imaging of [Ca^2+^]_i_ levels and AE**

In this study, both CD1 and Hybrid F1 mouse sperm were utilized. After undergoing capacitation, the sperm were centrifuged at 300g for 4 minutes. Subsequently, the cells were incubated in NC medium for 20 minutes with the addition of 1 μM Fluo-4 AM and 0.05% pluronic acid. After the incubation, the cells were centrifuged at 300g for 4 minutes and then resuspended in 200 μl of CAP medium. Recordings were conducted using a previously described method, with the recording medium (NC medium) containing FM4-64 concentrations ranging from 0.5 to 10 μM. For kymograph-like analysis, regions of interest (ROIs) were established throughout the midpiece during experiments performed at 10X magnification. [Ca^2+^]_i_ levels are represented as (F-F_0_)/F_0_ ratios after background subtraction, where F_0_ refers to the baseline determined by averaging the frames prior to stimulus application.

**Construction of a 3D kymograph**

The 3D kymograph was generated by merging the Fluo4 fluorescence kymograph (constructed in the same way as previously detailed for FM4-64) and the normalized diameter kymograph. Both kymographs possess three dimensions: time (x-axis), length along the midpiece (y-axis), and a color code to distinguish fluorescence intensity/diameter. By utilizing the plot_ly function of the R language environment, the 3D kymograph was plotted with time as the x-axis, length along the midpiece as the y-axis, and fluorescence intensity/diameter as the z-axis. While the color code for this dimension is not required, it remains present to facilitate visual interpretation of the data.

***Single-cell live imaging of* midpiece membrane and F-actin during AE**

Following swim-out, CD1 mouse sperm were incubated with 100 nM SiR-Actin for 10 min in NC medium. To maintain a constant signal and prevent the probe from interfering with actin dynamics, the concentration of SiR-Actin remained at 100 nM throughout the entire experiment. Recordings were performed as described earlier, using a recording medium (NC medium) containing 0.5 μM FM4-64 and 100 nM SiR-Actin.

**Fluorescence fluctuations super-resolution microscopy**

In this study, we employed fluorescence fluctuation super-resolution microscopy to process FM4-64 and SiR-actin images. The super-resolution radial fluctuations (SRRF) approach was utilized, leveraging the NanoJ plug-in within FIJI/ImageJ software (*17*, *41*). Each image consisted of 100 temporal frames, acquired with an exposure time of 10 msec per frame. During the analysis, specific parameters were set as follows: ring radius at 0.5, radiality magnification at 5, axes in ring at 8, and batch processing enabled. All other parameters were maintained at their default values.

Additionally, Memglow 700, Bodipy-GM1, and FM1-43 images were processed using the mean-shift super-resolution (MSSR) algorithm, facilitated by the MSSR plug-in within FIJI/ImageJ (*18*). Similar to the SRRF approach, each image was composed of 100 temporal frames, with an exposure time of 10 msec per frame. For the analysis, the following parameters were applied: AMP set to 5, PSF set to 10, order set to 0, and the interpolation type designated as bicubic.

**Colocalization Analysis using Manders' Coefficients**

To quantitatively evaluate the spatial overlap and potential associations between the fluorescently labeled biological structures, a colocalization analysis was conducted using Manders' colocalization coefficients (*23*). This analysis focused on the colocalization of FM4-64 and SiR-actin signals within the region of the midpiece of the flagellum, where the contraction initiation was observed.

Manders' colocalization coefficients (M1 and M2) were calculated using the R programming language. These coefficients provide a quantitative measure of colocalization between two fluorophores, with values ranging from 0 (indicating no colocalization) to 1 (representing complete colocalization). M1 corresponds to the fraction of intensity in channel 1 that overlaps with the intensity in channel 2, while M2 denotes the fraction of intensity in channel 2 that overlaps with the intensity in channel 1.

The utilization of Manders' coefficients offers advantages such as decreased sensitivity to variations in signal intensities and increased robustness against changes in background noise, as compared to other colocalization measurements, including Pearson's correlation coefficient and overlap coefficients. By determining the Manders' colocalization coefficients for the super-resolution images of the region of the flagellum's midpiece where the contraction initiation was observed, we objectively assessed the extent of spatial overlap between FM4-64 and SiR-actin. This analysis yielded valuable insights into potential interactions or associations between these two structures, ultimately contributing to an enhanced understanding of the underlying biological processes.

**Analysis of fluorescence peak position changes to assess relative proximity between plasma membrane and actin cytoskeleton**

The aim of this study was to assess the relative proximity between the plasma membrane (labeled with FM4-64) and the actin cytoskeleton (labeled with SiR-Actin) by analyzing the position changes of fluorescence peaks over time. Super-resolution radial fluctuations (SRRF) images were first generated for both structures. Subsequently, kymographs were computed from these SRRF images, and the spatial derivatives of the kymographs were calculated to identify the fluorescence peaks.

The R programming language was utilized for data analysis, computing the spatial derivatives of the kymographs to track the positions of the peaks. The fluorescence peak was identified at the location where the derivative transitioned between positive and negative values. The center of the cell was determined by averaging the positions of both membrane peaks, and subsequently, it was normalized to 0. Given the symmetrical nature of the fluorescence peaks around the cell center, the analysis focused on the right side of the graph (right peaks for FM4-64 and SiR-actin).

The distances to the peaks were plotted over time, followed by a linear regression. By examining the relative positions between the fluorescence maxima and their changes over time, the goal was to gain insights into the spatial relationship and potential interactions between the plasma membrane and the actin cytoskeleton.

**Three-dimensional stochastic optical reconstruction microscopy (3D-STORM)**

Following incubation under appropriate conditions (NC with DMSO or CAP with 100 µM progesterone), cells were washed with NC medium via centrifugation (4 min at 400g) and resuspended in NC medium. Sperm were seeded onto poly-L-lysine coated coverslips (Corning #1.5), air-dried for 10 min, fixed, and permeabilized with 0.3% fresh glutaraldehyde and 0.25% Triton-X100 in cytoskeleton buffer (CB, containing 10 mM MES, 150 mM NaCl, 5 mM EGTA, 5 mM glucose, and 5 mM MgCl_2_, pH = 6.1) for 1 min at room temperature. This was followed by three washes with CB (5 min at room temperature). Next, cells were incubated with 2% glutaraldehyde in CB for 15 min at room temperature and washed twice with CB (10 min at room temperature). Samples were treated with 0.1% NaBH_4_ (freshly prepared in PBS) for 7 min at room temperature to reduce background fluorescence, washed twice with PBS (5 min at room temperature), and incubated for 1 h at room temperature with Alexa 568-Peanut agglutinin (PNA, 0.01 mg/ml) and Alexa 647-Phalloidin (1:13) diluted in PBS. Sperm were then washed with PBS for 5 min and immediately mounted in STORM imaging buffer (50 mM Tris-HCl pH 8, 10 mM NaCl, 0.56 mg/ml glucose oxidase, 34 μg/ml catalase, 10% glucose, and 1% β-mercaptoethanol). Nonspecific staining was determined by incubating sperm in the absence of phalloidin (*13*).

Images were acquired using Andor IQ 2.3 software on a custom-built microscope equipped with an Olympus PlanApo 100x NA 1.45 objective and a CRISP ASI autofocus system (*42*, *43*). Alexa Fluor 647 was excited with a 642 nm laser (DL640-150-O, CrystaLaser, Reno, NV) under continuous illumination. Initially, the photo-switching rate was sufficient to provide a substantial fluorophore density. However, as fluorophores irreversibly photo-bleached, a 405 nm laser was introduced to enhance photo-switching. The intensity of the 405 nm laser was adjusted within the range of 0.01–0.5 mW to maintain an appropriate density of active fluorophores. Axial localization was achieved through astigmatism using a MicAO 3DSR adaptive optics system (Imagine Optic, Orsay, France), which allowed for both the correction of spherical aberrations and the introduction of astigmatism (*44*). Axial localization with adaptive optics enabled 3D reconstruction over a thickness of 1 μm (*13*). A calibration curve for axial localization was generated with 100 nm TetraSpeck microspheres (Invitrogen) immobilized on a coverslip (*45*). Images were acquired using a water-cooled, back-illuminated EMCCD camera (Andor iXon DU-888) operated at -85°C at a rate of 23 frames/s. A total of 50,000 frames were collected to generate each super-resolution image. Single-molecule localization, drift correction using image cross-correlation, and reconstruction were performed with ThunderSTORM (*46*). To analyze the molecular radial distributions, regions of interest (ROIs) of the flagellum were selected, ensuring they were in a straight line. The center of the flagella cross-section was initially identified by Gaussian fitting of the localization histograms along the x and y axes. Subsequently, the coordinates of the localized molecules were transformed into cylindrical coordinates to obtain the radial position (r) and azimuthal angle (Ɵ) (*13*, *47*, *48*). This approach allowed for a detailed examination of the spatial organization and molecular distribution of the sperm cell structures.

**Number and brightness**

To investigate structural rearrangements of the actin cytoskeleton during AE, a moment-based analysis was performed (*24*). This approach involved processing fluorescence signal intensity fluctuations to gain insight into actin filament polymerization and redistribution in the midpiece. Timelapse images of live sperm stained with SiR-actin during AE induction were captured, with 100 frames acquired every 30 seconds. For each consecutive 100-frame segment of the timelapse, temporal signal mean and variance were calculated, and number and brightness maps, as well as mean intensity sequences, were obtained (Figure S8A).

Prior to this step, a sCMOS sensor noise-correction algorithm (*25*) was applied to the images to reduce the effect of local digital noise variation on the analysis. Camera calibration maps were obtained as described elsewhere (*26*). Extended spatial resolution micrographs of the timelapses were obtained using MSSR, which allowed for the identification of nanoscale local enrichments of polymerized actin in the midpiece. Locations of these enrichments were selected and stored using the ROI tool in FIJI, with obtained ROIs consisting of 1 µm² areas over the identified local SiR-actin signal enrichment zones. By using the 'RImajeJROI' package for RStudio, saved ROIs were imported into the R environment, where they were utilized to extract information on the dynamic changes of actin signal as a consequence of AE induction (Figure S8B). This analysis provided valuable insights into the structural organization and dynamics of the actin cytoskeleton during the AE process.

**IVF assay**

Eight to 12-week-old F1 female mice were superovulated using equine chorionic gonadotropin (5 IU, PMSG; Syntex, Argentina) administered at 18:30, followed by human chorionic gonadotropin (5 IU, hCG; Syntex, Argentina) intraperitoneal injection 48 hours later. Cumulus oocyte complexes were collected from oviducts 12–13 hours post-hCG administration and placed in TYH IVF medium (which contains 25 mM NaHCO_3_ and 4 mg/ml BSA, without HEPES addition). The complexes were then inseminated with capacitated sperm at a final concentration ranging between 1x10^6^ and 2x10^6^ sperm/ml (a high concentration to allow polyspermy).

Following a 4-hour coincubation period at 37°C with 5% CO_2_, oocytes were stained with Hoechst 33342 (10 µg/ml) for 15 minutes and subsequently washed three times before being mounted on a glass slide. Oocytes were placed on 5 µl of CAP 0.5X modified TYH medium (7.5 mM NaHCO_3_ and 2.5 mg/ml BSA, 10 mM HEPES) containing 10 µM FM4-64, gently compressed beneath an 18 x 18 mm² coverslip supported by four solid Vaseline spots, and sealed with nail polish. This IVF assay protocol facilitated the study of fertilization events and sperm-oocyte interactions in a controlled in vitro environment.

**Sperm-oocyte fusion assay**

The sperm-oocyte fusion event was assessed using the zona-free oocyte penetration assay in conjunction with the Hoechst dye transfer technique (*28*–*30*). Cumulus oocyte complexes were collected as previously described. Oocytes were freed from cumulus cells by treatment with 0.2 mg/ml hyaluronidase and had their ZP removed by exposure to acid Tyrode's solution (pH 2.5) for 10-20 seconds (*49*).

Zona-free mouse oocytes were preloaded with 1 µg/ml Hoechst 33342 in TYH IVF medium for 5 minutes at 37°C. Following incubation, oocytes were washed three times for 20 minutes each in fresh TYH IVF medium and mounted in an oocyte-holding Petri dish (*27*) (MatTek glass bottom dishes, 35 mm Petri dish, 14 mm Microwell poly-d-lysine coated P35GC-1.5-14-C). Zona-free oocytes were placed on 5 µl of TYH IVF medium containing 10 µM FM4-64 and gently compressed under a 9 x 9 mm² coverslip supported by four solid Vaseline spots. TYH IVF medium containing 10 µM FM4-64 was added until reaching a final volume of 200 μl, and the medium was then covered with prewarmed (37°C) mineral oil (M8410, Sigma).

The dish was placed into the microscope incubation chamber at 37°C with 5% CO_2_, and oocytes were inseminated with capacitated sperm at a final concentration of 1x10^5^ cells/ml. For Figure 7, EGFP-DsRed2 sperm were used, while for Figure 8, F1 sperm loaded with Fluo4 (as described above) were utilized. This assay allowed for the investigation of sperm-oocyte fusion events and the assessment of the transfer of Hoechst dye between the two cell types.

**Statistical analysis**

Data are presented as mean ± standard error of the mean (SEM) from a minimum of three independent experiments for all measurements. Statistical analyses were conducted using GraphPad Prism v4.0 (San Diego, CA, USA) or the R language environment [R 3.3.3 GUI 1.69 Mavericks build (7328)]. The specific statistical analysis employed is indicated in the relevant figure legend. A p-value of less than 0.05 was considered statistically significant.
