## Supplementary Table 1 for "Reorganization of the Flagellum Scaffolding Induces a Sperm Standstill During Fertilization"

**Table S1: Summary of samples, microscopes, imaging conditions and probes used**. The notation DsRed2* refers to the co-excitation of DsRed2 (from the mitochondria of the EGFP/DsRed2 transgenic model) with FM4-64, see Figure S1.

| **Figure** | **Sample** | **Microscope** | **Imaging conditions** | **Probes** |
| --- | --- | --- | --- | --- |
| 1A | Transgenic EGFP-DsRed2 sperm | Fluorescence spinning disk confocal microscope (Olympus IX83-DSU) and a 10X objective (UPLSAPO, NA 0.4) with a sCMOS camera (Andor, Zyla). | Exposure time: 500 msec for both EGFP and DsRed2.  Acquisition frequency: 1 image every 1 min, for 15 min.  Filters: for EGFP, the filter U-FBWA Ex 460-495 nm /Em 510-550 nm; for DsRed2 the filter U-FGWA Ex 530-550 nm/Em 575-625 nm. | - |
| 1B | Transgenic EGFP-DsRed2 sperm | Eclipse TE 300 Nikon microscope with a 40X oil immersion objective (PlanApo N, NA 1.42) with EMCCD camera (Andor Technology, Belfast, UK). | Exposure time: 500 msec.  Acquisition frequency: 2 images every 1 sec for 5 min.  Filters: Led cyan (3.15 A, Luminus Devices, Woburn, MA) Bandpass, excitation filter (HQ 480 nm /40X), dichroic mirror (Q505lp), and emission filter (HQ 535 nm /50M, Chroma Technology, Bellows Falls, VT). | - |
| 1D | Transgenic EGFP-DsRed2 sperm | Fluorescence spinning disk confocal microscope (Olympus IX83-DSU) and a 10X objective (UPLSAPO, NA 0.4) with a sCMOS camera (Andor, Zyla). | Exposure time: 500 msec for both EGFP and FM4-64.  Acquisition frequency: 1 image every 1 min, for 15 min.  Filters: for EGFP, the filter U-FBWA Ex 460-495 nm /Em 510-550 nm; for FM4-64 the filter ETCY5 Ex 590-650 nm/Em 665-735 nm. | FM4-64 10 µM |
| 1E, 1F and 1G, left panel | Transgenic EGFP-DsRed2 sperm | Nikon (Melville, NY,USA)-inverted microscope (Eclipse Ti-U) and a 60X oil immersion objective (plan Apo TIRF DIC, NA 1.45, Nikon) with an Andor Ixon 3 EMCCD camera model DU-8970-C00#B (Andor Technology, Belfast UK) or Hammamatsu Orca-100 CCD model C4742-95 (Hammamatsu Photonics, Bridgewater, NJ USA) | Exposure time: 100 msec for FM4-64, 350 msec for EGFP.  Acquisition frequency: 1 image every 2 sec, for 5 min.  Filters: for EGFP, filter set GFP 96343, dichroic mirror (D): 495 nm, Ex 470 nm/40, barrier 525 nm/50 (Nikon); for FM4-64, the filter set Wide Green 11007v2, D: 565 nm dcxt, Ex 535 nm/50, Em 590 nm lpv2 (Chroma Technology Corporation) | FM4-64 10 µM |
| 1E, 1F and 1G, right panel | Transgenic EGFP-DsRed2 sperm | Fluorescence spinning disk confocal microscope (Olympus IX83-DSU) and a 60X oil immersion objective (Plan Apo, NA 1.4) with a sCMOS camera (Andor, Zyla). | Exposure time: 500 msec for both EGFP and FM4-64.  Filters: for EGFP, the filter U-FBWA Ex 460-495 nm/Em 510-550 nm; for FM4-64 the filter ETCY5 Ex 590-650 nm/Em 665-735 nm. | FM4-64 10 µM |
| 2A, 2B | CD1 sperm | NanoImager S microscope (Oxford Nanoimaging Ltd), equipped with a 100X oil-immersion objective (NA 1.4, Olympus) and a sCMOS camera (Hamamatsu ORCA-Flash4.0 V3 Digital CMOS camera) | Exposure time: 10 msec.  Acquisition frequency: 100 images every 30 sec for 20 min.  Filters: for excitation a 561 nm laser was used, the emission was collected in channel 1 (665-705 nm) | FM4-64 0.5 μM |
| 2G | Transgenic EGFP-DsRed2 sperm | Zeiss LSM880 microscope equipped with an 63X oil-immersion objective (NA 1.4) and a EMCCD Andor iXon 897 camera. | Laser power: for DsRed 5%, for EGFP 1.2% and for Alexa Fluor 647 24%. | ɑ-GLUT3 1:100, Alexa Fluor 647 1:200 |

| 4A, 4B, 4C, 4D and S3 | Hybrid F1 (C57BL/6 × Balb/C) sperm | Fluorescence spinning disk confocal microscope (Olympus IX83-DSU) and a 10X objective (UPLSAPO, NA 0.4) with a sCMOS camera (Andor, Zyla). | Exposure time: 500-200 msec for both Fluo4 and FM4-64.  Acquisition frequency: 1 image every 10 sec, for 9 min.  Filters: for Fluo4, the filter U-FBWA Ex 460-495 nm/Em 510-550 nm; for FM4-64 the filter ETCY5 Ex 590-650 nm/Em 665-735 nm. | FM4-64 10 µM, Fluo4 1 µM |
| --- | --- | --- | --- | --- |
| 4C | CD1 sperm | NanoImager S microscope (Oxford Nanoimaging Ltd), equipped with an 100X oil-immersion objective (NA 1.4, Olympus) and a sCMOS camera (Hamamatsu ORCA-Flash4.0 V3 Digital CMOS camera) | Exposure time: 10 msec.  Acquisition frequency: 100 images every 30 sec for 20 min.  Filters: for excitation of Fluo4 a 473 nm laser was used, the emission was collected in channel 0 (band 1: 498-551 nm); for excitation of FM4-64 a 561 nm laser was used, the emission was collected in channel 1 (665-705 nm) | FM4-64 0.5 µM, Fluo4 1 µM |
| 5A, 5B and S5 | CD1 sperm | NanoImager S microscope (Oxford Nanoimaging Ltd), equipped with an 100X oil-immersion objective (NA 1.4, Olympus) and a sCMOS camera (Hamamatsu ORCA-Flash4.0 V3 Digital CMOS camera) | Exposure time: 10 msec.  Acquisition frequency: 100 images every 30 sec for 20 min.  Filters: for excitation of FM4-64 a 561 nm laser was used, the emission was collected in channel 1 (665-705 nm); for excitation of SiR-actin a 640 nm laser was used, the emission was collected in channel 1 (665-705 nm) | FM4-64 0.5 µM, SiR-actin 0.1 µM |
| 5D, 5E, 5F and 5G | CD1 sperm | See above | | |
| 6 | Transgenic EGFP-DsRed2 sperm | Fluorescence spinning disk confocal microscope (Olympus IX83-DSU) and a 20X (UPLSAPO, NA 0.75) or 40X magnification objective (UPLSAPO, NA 0.95) with a sCMOS camera (Andor, Zyla). | Exposure time: 10 msec for DIC, 50 msec for Hoechst, 120 ms for EGFP and DsRed2* and 500 msec for FM4-64.  Acquisition frequency: 1 image every 5 µm with a total of 10-15 planes.  Filters: for Hoechst 33342, the filter LF405-B Ex 368-415 nm/Em 426-477 nm; for EGFP, the filter U-FBWA Ex 460-495 nm/Em 510-550 nm; for DsRed2*, the filter U-FGWA Ex 530-550 nm /Em 575-625 nm and for FM4-64 the filter ETCY5 Ex 590-650 nm/Em 665-735 nm. | FM4-64 10 µM, Hoechst 33342 10 µg/ml |
| 7, S7 | Transgenic EGFP-DsRed2 sperm | Fluorescence spinning disk confocal microscope (Olympus IX83-DSU) and a 10X (UPLSAPO, NA 0.4) or 20X magnification objective (UPLSAPO, NA 0.75) with a sCMOS camera (Andor, Zyla). | Exposure time: 10 msec for DIC, 50 msec for Hoechst, 120 msec for EGFP and DsRed2-FM4-64 and 500 msec for FM4-64.  Acquisition frequency: 1 image every 1 min for 60-120 min and 1 image every 7 µm with a total of 10-15 planes.  Filters: for Hoechst 33342, the filter LF405-B Ex 368-415 nm/Em 426-477 nm; for EGFP, the filter U-FBWA Ex 460-495 nm/Em 510-550 nm; for DsRed2*, the filter U-FGWA Ex 530-550 nm /Em 575-625 nm and for FM4-64 the filter ETCY5 Ex 590-650 nm/Em 665-735 nm. | FM4-64 10 µM, Hoechst 33342 1 µg/ml |
| 8 | Hybrid F1 (C57BL/6 × Balb/C) sperm | Fluorescence spinning disk confocal microscope (Olympus IX83-DSU) and a 20X magnification objective (UPLSAPO, NA 0.75) with a sCMOS camera (Andor, Zyla). | Exposure time: 10 msec for DIC, 40 msec for Hoechst, 300 msec for Fluo4 and 100 msec for FM4-64.  Acquisition frequency: 1 image every 1 min for 60-120 min and 1 image every 7 µm with a total of 10-15 planes.  Filters: for Hoechst 33342, the filter LF405-B Ex 368-415 nm /Em 426-477 nm; for Fluo4, the filter U-FBWA Ex 460-495 nm /Em 510-550 nm and for FM4-64, the filter U-FGWA Ex 530-550 nm /Em 575-625 nm. | FM4-64 10 µM, Hoechst 33342 1 µg/ml, Fluo4 1 µM |
| S1 | Transgenic EGFP-DsRed2 sperm | Fluorescence spinning disk confocal microscope (Olympus IX83-DSU) and a 10X (UPLSAPO, NA 0.4) or 20X magnification objective (UPLSAPO, NA 0.75) with a sCMOS camera (Andor, Zyla). | Exposure time: 10 msec for DIC, 120 msec for EGFP and DsRed2* and 500 msec for FM4-64.  Acquisition frequency: 1 image every 10 msec for DIC, 1 image every 120 msec for EGFP and DsRed2* and 1 image every 500 msec for FM4-64 for 3 min every 5 min.  Filters: for EGFP, the filter U-FBWA Ex 460-495 nm /Em 510-550 nm; for DsRed2*, the filter U-FGWA Ex 530-550 nm /Em 575-625 nm and for FM4-64 the filter ETCY5 Ex 590-650 nm/Em 665-735 nm. | FM4-64 10 µM |
| S2A | Transgenic EGFP-DsRed2 sperm | NanoImager S microscope (Oxford Nanoimaging Ltd), equipped with an 100X oil-immersion objective (NA 1.4, Olympus) and a sCMOS camera (Hamamatsu ORCA-Flash4.0 V3 Digital CMOS camera) | Exposure time: 10 ms.  Acquisition frequency: 100 images every 10 ms.  Filters: for excitation of Memglow-700 a 640 nm laser was used, the emission was collected in channel 1 (665-705 nm), for excitation of EGFP a 473 nm laser was used, the emission was collected in channel 0 (band 1: 498-551 nm) | Memglow-700 0.2 µM |
| S2B | Transgenic EGFP-DsRed2 sperm | NanoImager S microscope (Oxford Nanoimaging Ltd), equipped with an 100X oil-immersion objective (NA 1.4, Olympus) and a sCMOS camera (Hamamatsu ORCA-Flash4.0 V3 Digital CMOS camera) | Exposure time: 10 msec.  Acquisition frequency: 100 images every 10 msec.  Filters: for excitation a 473 nm laser was used, the emission was collected in channel 0 (band 1: 498-551 nm) | Bodipy-GM1 0.1 mg/mL |
| S2C | Transgenic EGFP-DsRed2 sperm | NanoImager S microscope (Oxford Nanoimaging Ltd), equipped with an 100X oil-immersion objective (NA 1.4, Olympus) and a sCMOS camera (Hamamatsu ORCA-Flash4.0 V3 Digital CMOS camera) | Exposure time: 10 msec.  Acquisition frequency: 100 images every 10 msec.  Filters: for excitation a 473 nm laser was used, the emission was collected in channel 0 (band 1: 498-551 nm) | FM1-43 1 µM |
| S4 | Transgenic EGFP-DsRed2 sperm | NanoImager S microscope (Oxford Nanoimaging Ltd), equipped with an 100X oil-immersion objective (NA 1.4, Olympus) and a sCMOS camera (Hamamatsu ORCA-Flash4.0 V3 Digital CMOS camera) | Exposure time: 10 msec.  Acquisition frequency: 100 images every 10 msec.  Filters: for excitation of SiR-actin a 640 nm laser was used, the emission was collected in channel 1 (665-705 nm),; for excitation of Bodipy-GM1/EGFP a 473 nm laser was used, the emission was collected in channel 0 (band 1: 498-551 nm) and for excitation of DsRed2 a 561 nm laser was used, the emission was collected in channel 1 (665-705 nm). | Bodipy-GM1 0.1 mg/mL, SiR-actin 0.1 µM |
