## Supplementary Figures and movies legends for "Reorganization of the Flagellum Scaffolding Induces a Sperm Standstill During Fertilization"

**Supplementary Figures and videos legends.**

**Figure S1. Gradual decrease in flagellar beat frequency following acrosomal exocytosis.** Motility analysis was conducted for capacitated EGFP-DsRed2 sperm. Cells were immobilized on laminin-coated coverslips and incubated in a recording medium (rec) containing 10 μM FM4-64. A) Due to the spectral characteristics of DsRed2 (in the transgenic model), FM4-64, and the microscope configuration, both DsRed2 and FM4-64 emissions were observed using the U-FGWA excitation 530-550 nm / emission 575-625 nm filter (shown in yellow). In contrast, using the ETCY5 excitation 590-650 nm / emission 665-735 nm filter allowed for the visualization of only FM4-64 emission (in red). As a result, the images display the plasma membrane and mitochondria in yellow (with DsRed2* notation), while the plasma membrane appears in red. B-C) The figure presents a representative time series of sperm without AE (B) and progesterone-induced AE (C, prog, 100 μM). The upper panel displays DIC images processed by a cell tracking system to measure flagellar beat frequency, along with EGFP, DsRed2*, and FM4-64 images. Scale bar = 10 μm. The lower graphs illustrate the frequency analysis (right y-axis) for control sperm (with the addition of rec, indicated by an arrowhead) and progesterone-induced AE. The percentage of immotile cells is shown on the left y-axis. These images represent at least five independent experiments, with 11 cells analyzed for the control (rec) group and 22 cells for the progesterone-induced AE group. D) Relationship between cell death and FM4-64 fluorescence capacitated sperm stimulated with ionomycin. Image-based flow cytometry analysis of non-capacitated mouse sperm loaded with FM4-64 and Sytox Blue dyes, with one and two minutes of incubation time, respectively. The quadrants in the left panel show: Sytox Blue+ / FM4-64 low (4.52%), Sytox Blue+ / FM4-64 high (60.6%), Sytox Blue- / FM4-64 low (20.5%), and Sytox Blue- / FM4-64 high (13.5%). Each quadrant indicates the percentage of the total sperm population exhibiting the corresponding staining pattern. Axes are presented on a log10 scale of arbitrary units of fluorescence. The right panel shows representative single-cell images corresponding to the four categorized sperm populations from the flow cytometry analysis in the left panel.

**Figure S2. Correlation between FM4-64 increase and midpiece contraction**. A-C)	Representative MSSR-processed images of acrosome-intact (AI) and acrosome-reacted (AR) EGFP-DsRed2 transgenic sperm stained with membrane dyes Memglow 700 (A, violet), Bodipy-GM1 (B), and FM1-43 (C). Insets in the sperm head images reveal AE in wide-field views: in A), AE is visible with EGFP in green and the membrane in violet, while in B) and C), the acrosome has the same color as the membrane. Scale bar = 1 μm. D) C	orrelation between FM4-64 midpiece fluorescence (y-axis) and normalized diameter (x-axis), with data normalized to the mean of the frames prior to AE induction. E-H) R	epresentative contraction kymographs and diameter measurements for ionomycin-induced AE (10 μM, E-F) and spontaneous AE (G-H), respectively, with one or multiple contraction foci. In the contraction kymographs, yellow lines indicate the separation between midpiece sections, and colored spots mark the locations where super-resolution kymographs were generated. A dotted vertical line in both kymograph and diameter measurement graphs signifies the point of induction. The data presented are derived from at least five independent experiments, with 36 cells analyzed. For E, horizontal scale bar = 5 min and vertical scale bar = 2 μm; for F, horizontal scale bar = 4 min and vertical scale bar = 2 μm; for G, horizontal scale bar = 5 min and vertical scale bar = 1 μm and for H, horizontal scale bar = 5 min and vertical scale bar = 2 μm.

**Figure S3. [Ca^2+^]_i_ concentration drives midpiece contraction in Ionomycin-stimulated sperm.** A) The representative time series demonstrates [Ca^2+^]_i_ and AE dynamics. Capacitated F1 sperm, loaded with Fluo4 AM, were immobilized on concanavalin A-coated coverslips and incubated in a recording medium containing 10 μM FM4-64. Ionomycin (iono, 10 μM) was added as indicated by arrowheads. Beneath each frame in the Fluo4 (green) and FM4-64 (red) images, a color code displays the normalized intensity of the fluorescence signal (scale bar on the right of the panel). Scale bar = 10 μm. B) The kymograph-like analysis of 20 ionomycin-stimulated cells shows each row depicting the [Ca^2+^]_i_ (upper) and membrane (lower) dynamics of a single cell over time. A white dotted line indicates the moment of ionophore addition. The images presented are representative of at least five independent experiments.

**Figure S4. Unaltered distance between F-actin and mitochondrial network after AE indicates consistent mitochondrial network organization in EGFP-DsRed2 Transgenic Sperm.** A) Representative super-resolution images of the plasma membrane (Bodipy-GM1), actin cytoskeleton (SiR-actin), and mitochondria (DsRed2) for acrosome-intact (upper panel) and acrosome-reacted (lower panel) EGFP-DsRed2 transgenic sperm. Insets in the Bodipy-GM1 images display the sperm head with AE in wide-field images. Scale bar = 1 μm. The left panel illustrates the normalized fluorescence intensity for all three dyes.

**Figure S5. Plasma membrane approaches actin cytoskeleton during midpiece contraction.** A-C	) Super-resolution kymographs display FM4-64 (upper) and SiR-actin (lower) dynamics, computed at the proximal region of the midpiece. A dotted white line indicates the point of	induction for ionomycin-induced AE (10 μM, A), progesterone-induced AE (100 μM, B) and spontaneous AE (C) . D-F	) Normalized fluorescence of FM4-64 (red) and SiR-actin (blue) across the midpiece of the flagellum is shown over time for ionomycin-induced AE (10 μM, D), progesterone-induced AE (100 μM, E) and spontaneous AE (F). A horizontal black bar represents the initial distance between FM4-64 peaks. G	) Using the data from panels A and D	, this panel presents an analysis of the changes in the positions of fluorescence peaks, measured as the distance to the center of the flagella, over time. FM4-64 peaks are shown in red, and SiR-actin peaks are shown in blue. The numbers near the lines indicate the slopes, represented in μm/min⁻¹. H	) Slope values for no reaction, spontaneous, progesterone, and ionomycin-induced AE for FM4-64 and SiR-actin. Data are presented as mean ± SEM, and the images shown are representative of at least five independent experiments, with 36 cells analyzed. I	) The distance between the plasma membrane and actin double helix in the midpiece for acrosome-intact and reacted sperm is analyzed and represented as box plots. A box plot is a graphical representation of data that displays the median, quartiles, and a summary of the data distribution. Wilcoxon test was performed, ****p<000.1.

**Figure S6. Actin cytoskeleton reorganization in the midpiece during AE.** A) Changes in SiR-actin fluorescence signal intensity (int), Number (N), and Brightness (B) following AE induction. After 2-3 minutes of time-lapse recording, progesterone (prog, 100 μm) or ionomycin (iono, 10 μm) was added to the media. Using the generated ROIs for each cell (Figure S8B), SiR-actin signal was measured at each of these regions before and after induction (at its maximum increase point). The reported values indicate the maximum times-fold increase in SiR-actin int/N/B. The sample size n of this analysis represents the total number of ROIs analyzed across all cells for each condition: negative control (non-reacted sperm): n = 72, cells = 5, mice = 2; progesterone: n = 315, cells = 28, mice = 11; ionomycin: n = 324, cells = 22, mice = 10; spontaneous: n = 184, cells = 14, mice = 5. Statistical significance assessed by non-parametric Wilcoxon test. B) Proposed model of the structural rearrangements of the actin cytoskeleton in the midpiece during AE: Based on the results obtained from the moment-based analysis shown in A), this work suggests that the actin filament network in the midpiece experiences dynamic polymerization, evidenced by the increase in SiR-actin fluorescence and Number. Additionally, reorganization is indicated by the overall increase in the magnitude and dispersion of the brightness measurement, as a consequence of the AE.

**Figure S7. Midpiece contraction occurs following sperm-egg fusion.** A) DIC image of a sperm-oocyte interaction, with the area depicted in higher magnification in B. Scale bar = 20 μm. B) Representative time series of sperm-oocyte fusion assay experiments using EGFP-DsRed2 transgenic sperm. Oocytes were stained with 1 μg/ml Hoechst and 10 μM FM4-64. DIC, Hoechst, EGFP, DsRed2*, and FM4-64 images are shown over time. Scale bar = 10 μm. The images shown are representative of at least four independent experiments.

**Figure S8. Moment-based Analysis Overview.** A) The diagram illustrates the AE induction time-lapse recording process. Imaging was performed using an ONI Nanoimager-S microscope, featuring a 100x 1.49 NA oil-immersion objective. Excitation of FM4-64 and SiR-actin was achieved using 561 nm and 640 nm lasers, respectively. Scale bar = 2 μm.B) Time-lapse recordings from both FM4-64 (red) and SiR-actin (cyan) channels were employed to generate an extended-resolution image reconstruction. Scale bar = 2 μm. This allowed for the detection of nanoscale localized enrichments of polymerized actin. Number and Brightness analysis was executed using a custom RStudio code. The following MSSR parameters were applied: AMP = 3; FWHM of PSF = 2.44; Order = 0; Interpolation = bicubic; Temporal analysis = Mean (100 frames).

**Supplementary movies legends**

**Movie S1.** Representative movie of transgenic EGFP-DsRed2 sperm attached to concanavalin A-coated coverslips, with AE induced by 100 μM progesterone as indicated with Prog. DsRed2 is presented in orange, and EGFP in green. White squares indicate cells with spontaneous AE (prior to induction), while pink squares highlight cells with progesterone-induced AE. Related to Figure 1A.

**Movie S2.** Representative movie of transgenic EGFP-DsRed2 sperm that have already experienced AE, attached to laminin-coated coverslips. The left panel displays a cell with motility after AE, and the right panel shows an immotile cell. DsRed2 is presented in orange, and EGFP in green. Related to Figure 1B

**Movie S3.** Representative movie of transgenic EGFP-DsRed2 sperm stained with 10 μM FM4-64 and attached to concanavalin A-coated coverslips, with AE induced by 100 μM progesterone as indicated with Prog. FM4-64 is presented in red, and EGFP in green. White squares indicate cells exhibiting patterns I, II, or III after progesterone induction. Related to Figure 1D.

**Movie S4.** Representative movies of capacitated transgenic EGFP-DsRed2 sperm stained with 10 μM FM4-64. Left panel displays an acrosome-intact, motile sperm (Pattern I), middle panel shows an acrosome-reacted sperm with motility and low FM4-64 midpiece fluorescence (Pattern II), and right panel presents an acrosome-reacted sperm with no motility and high FM4-64 midpiece fluorescence (Pattern III). In all three cases, cells were induced with 100 μM progesterone. FM4-64 is presented in red, and EGFP in green. Related to Figure 1E, 1F and 1G.

**Movie S5.** Representative movie of capacitated CD1 sperm stained with 0.5 μM FM4-64 64 and attached to concanavalin A-coated coverslips with spontaneous AE. Following acquisition, images were analyzed using SRRF. FM4-64 is presented in red. Related to Figure 2B and 2C.

**Movie S6.** Representative movies showing [Ca^2+^]_i_ and AE dynamics for a control case (addition of recording medium, rec), progesterone (Prog, 100 μM) and ionomycin-induced AE (Iono, 10 μM). Capacitated F1 sperm, loaded with Fluo4 AM, were immobilized on concanavalin A-coated coverslips and incubated in a recording medium containing 10 μM FM4-64. Additions were added when indicated with Rec, Prog or Iono. Fluo4 is presented in green and FM4-64 in red. Related to Figures 4A, 4C and S3A.

Movie S7. Representative movie of IVF assay. DIC images are shown. Related to Figure 6A.

**Movie S8**	**.** Representative movie of sperm-oocyte fusion assay experiments using EGFP-DsRed2 sperm. Oocytes were stained with 1 μg/ml Hoechst and 10 μM FM4-64. DIC images are presented in grey scales, Hoechst in blue, EGFP is green, DsRed2* in yellow, and FM4-64 in red. Sperm of interest is indicated with a white arrow.

**Movie S9**	**.** Representative movie of sperm-oocyte fusion assay experiments using wild-type sperm loaded with 1 μM Fluo-4. Oocytes were stained with 1 μg/ml Hoechst and 10 μM FM4-64. DIC images are presented in grey scales, Hoechst in blue, Fluo4 in green, and FM4-64 in red.
